# NPEPPS is a novel and druggable driver of platinum resistance

**DOI:** 10.1101/2021.03.04.433676

**Authors:** Robert T. Jones, Mathijs Scholtes, Andrew Goodspeed, Maryam Akbarzadeh, Saswat Mohapatra, Lily Elizabeth Feldman, Hedvig Vekony, Annie Jean, Charlene B. Tilton, Michael V. Orman, Shahla Romal, Cailin Deiter, Tsung Wai Kan, Nathaniel Xander, Stephanie Araki, Molishree Joshi, Mahmood Javaid, Ryan Layer, Teemu D. Laajala, Sarah Parker, Tokameh Mahmoudi, Tahlita Zuiverloon, Dan Theodorescu, James C. Costello

## Abstract

There is an unmet need to improve efficacy of platinum-based cancer chemotherapy. Using multi-omic assessment of cisplatin-responsive and -resistant human bladder cancer cell lines and whole-genome CRISPR screens, we identified Puromycin-Sensitive Aminopeptidase, NPEPPS, as a novel driver of cisplatin resistance. NPEPPS depletion sensitizes resistant bladder cancer cells to cisplatin *in vitro* and *in vivo*. Conversely, overexpression of NPEPPS in sensitive cells increased cisplatin resistance. We show that NPEPPS affects treatment response by regulating intracellular cisplatin concentrations. Patient-derived organoids (PDOs) generated from bladder cancer samples before and after cisplatin-based treatment, and from patients who did not receive cisplatin, were evaluated for sensitivity to cisplatin and they were found to be concordant with clinical response. In PDOs, shRNA depletion or pharmacologic inhibition of NPEPPS led to increased cisplatin sensitivity, while NPEPPS overexpression had the opposite effect. Our data present NPEPPS as a novel and druggable driver of cisplatin resistance by regulating intracellular cisplatin concentrations, along with providing the preclinical data to support clinical trials combining NPEPPS inhibition with cisplatin.

## INTRODUCTION

Platinum-based chemotherapy has been successful in testicular, ovarian, bladder, head and neck, and lung cancer and remains a standard of care for many patients despite the advent of immunotherapy (1,2). However, dose-dependent side effects and chemoresistance have reduced the suitability and effectiveness of these drugs. The discovery of more effective agents or the development of strategies that improve efficacy of platinum-based regimens are approaches that can address these issues. Here we describe results from the latter approach, using bladder cancer (BCa) as a clinically relevant, and translationally tractable tumor models.

Bladder cancer accounts for 430,000 new diagnoses and 170,000 deaths worldwide annually (3). Cisplatin-based combination chemotherapy remains the first-line, standard of care for metastatic BCa, providing a 5-10% cure rate. However, up to 30% of patients are ineligible for cisplatin-based treatment (4) and are commonly offered immune checkpoint therapies (ICT) (5). However, ICT requires a PD-L1 diagnostic test, for which only ∼25% of patients meet eligibility (6). Cisplatin-based combination chemotherapy is also standard of care in the neoadjuvant (NAC) setting for the management of localized muscle-invasive BCa (MIBC) (7,8). However, NAC adoption has been slow due to cisplatin toxicity and eligibility, along with the relatively small survival benefit of 5-15% over immediate cystectomy (9). Importantly, in both the metastatic and NAC settings, patient selection and therapeutic efficacy remain critical unresolved challenges (10).

Large-scale efforts have performed whole genome loss-of-function screening across hundreds of cancer cell lines using CRISPR- and shRNA-based libraries to define pan-cancer and context-specific genetic dependencies (11–14). While these efforts defined genetic dependencies in cancer, a limitation is that cells were grown under basal growth conditions in the absence of any treatment. Those studies were also performed in cells that had not acquired resistance to any treatments. To better understand the functional drivers of therapeutic resistance in the context of platinum drug treatment, here we performed CRISPR screens in the presence and absence of the therapy of interest (15–18), and in cells that have acquired resistance to the treatment itself (**Figure 1A**). Complementing these results with multi-omic profiling of treatment-responsive and -resistant cells allowed us to prioritize gene candidates for the best potential future clinical utility. Using these data, which make available as a public resource https://bioinformatics.cuanschutz.edu/BLCA_GC_Omics/), we identified Puromycin-Sensitive Aminopeptidase, NPEPPS, as a novel driver of cisplatin resistance and validated this finding *in vitro* and *in vivo*. We then show that NPEPPS has its effect by directly regulating the intracellular concentrations of cisplatin. To determine the translational relevance of this discovery, tumor-derived organoids generated from patients before and after cisplatin-based chemotherapy, were used to validate and strengthen our results from bladder cancer cells and xenografts. In cells, xenografts, and organoids, we show that pharmacological inhibition of NPEPPS by tosedostat, a clinically used small molecule that inhibits NPEPPS, phenocopies genetic depletion of NPEPPS. Taken together our data supports NPEPPS as a therapeutic target and provides a compelling rationale for combining NPEPPS inhibition with cisplatin to improve bladder cancer patient outcomes.

**Figure 1.**
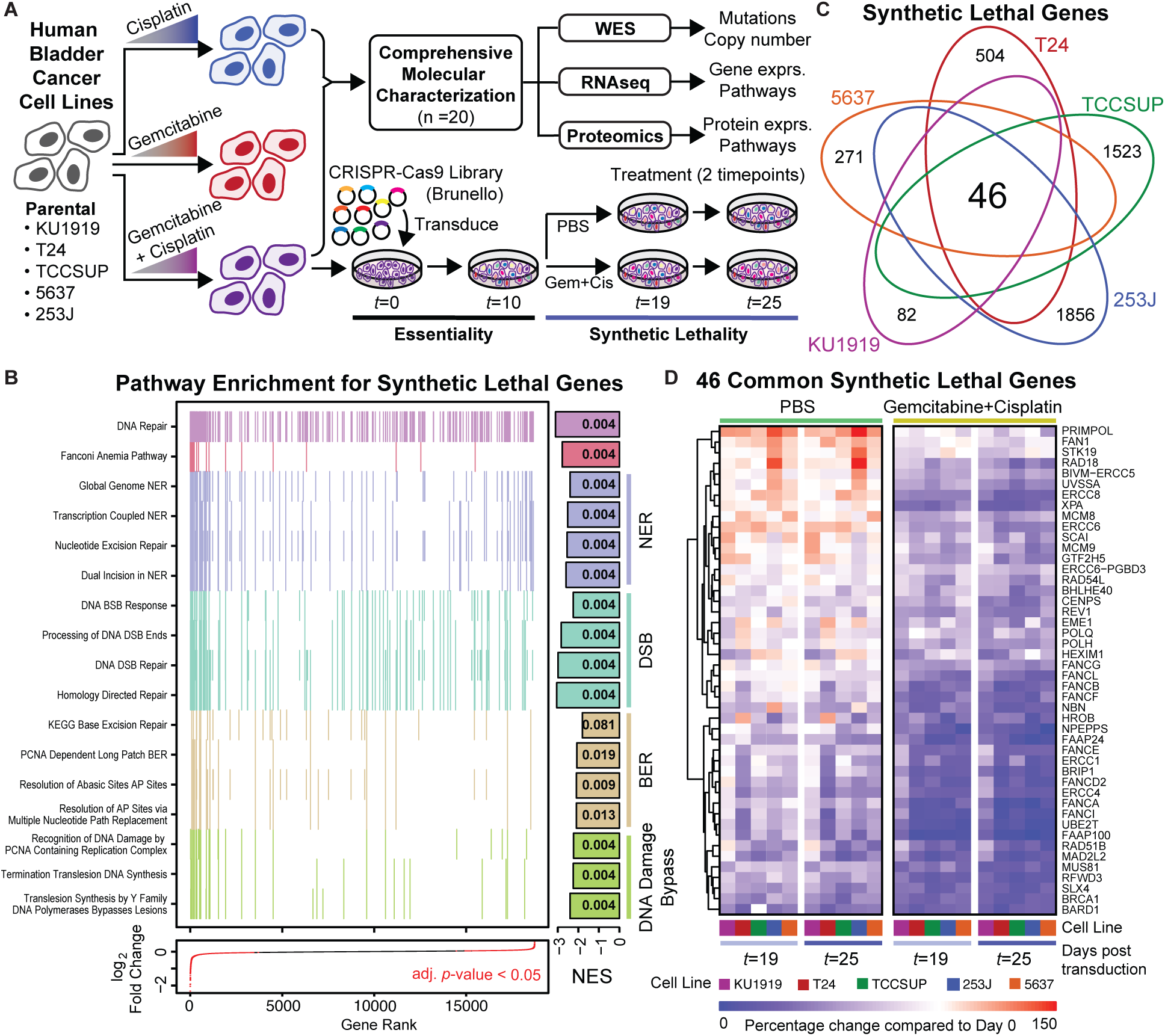
Project overview and synthetic lethal screen results. (**A**) Human bladder cancer cell lines were made resistant to cisplatin, gemcitabine, or gemcitabine plus cisplatin through dose escalation. All cell lines were profiled using -omic technologies. The gemcitabine plus cisplatin-resistant cells were subjected to a pooled CRISPR screen to identify synthetic lethal gene-to-drug relationships. (**B**) Aggregate gene set enrichment results for the synthetic lethal screen ranked by log_2_ fold change across all cell lines reveal DNA damage response and repair pathways. Each tick mark represents a gene in the associated pathway. The bars at the right are normalized enrichment scores (NES) with the FDR corrected p-values reported in the bars. (**C**) The intersection across the CRISPR screen results identified 46 common synthetic lethal genes; all counts and gene annotations are reported in **Figure S2**. (**D**) The percentage change in the aggregate of the sgRNAs targeting the 46 commonly synthetic lethal genes are reported across phosphate-buffered saline (PBS) or gemcitabine plus cisplatin treatment arms of the CRISPR screen. Cell lines are coded with the same colors throughout all figures.

## MATERIALS AND METHODS

### Cell Culture

All human BCa cell lines as reported in the Key Resource Table were obtained from the Resistant Cancer Cell Line (RCCL) Collection and were grown in Iscove’s Modified Dulbecco’s Medium (IMDM) with 10% Fetal Bovine Serum (FBS). Cells were passaged every two to three days. Resistance to gemcitabine and cisplatin was confirmed at the reported resistance doses from the RCCL (**Table S1** and **Figure S1**). Lentivirus production utilized 293FT cells (ThermoFisher), which were maintained in DMEM (high glucose) supplemented with 0.1mM non-essential amino acids (NEAA), 6mM L-glutamine, 1mM sodium pyruvate, and 500μg/mL geneticin (G418) with 10% FBS added. Cells were routinely monitored for mycoplasma and confirmed negative at multiple times during this study using MycoAlert (Lonza). All cells were grown at 37°C with 5% CO_2_ in a humidified incubator.

All molecular characterization efforts (RNA sequencing, whole exome sequencing, and mass spectrometric proteomics) were performed on cells from independent passages and in drug-free, complete media to identify stable molecular changes rather than treatment-induced transient responses. Cells were routinely passaged through drug-containing media at the resistant doses (**Table S1**) to confirm resistance was maintained and early passage cells were utilized whenever possible.

### RNA sequencing

#### Sample preparation

All cell lines were grown for several passages in the absence of antibiotics, gemcitabine or cisplatin. Cell pellets were snap frozen from sub-confluent dishes from 3 separate passages (replicates) for each of the 20 cell lines sequenced (5 cell lines, each with 4 derivatives: parental, G-resistant, C-resistant, GC-resistant). RNA was extracted using the RNAeasy Plus Kit (Qiagen). Cells were lysed and passed through QIAShredder column (Qiagen) according to the manufacturer’s protocol. gDNA elimination columns (Qiagen) were used to remove any residual gDNA from the purified RNA. RNA integrity was assessed on the High Sensitivity ScreenTape Assay on the Tape Station2200 (Agilent) and only samples with an RIN score of 8 or higher were used for sequencing. RNA library preparation was performed using the Universal Plus mRNA –Seq +UDI kit (Nugen) according to the manufacturer’s specification. Each library was sequenced to a minimum of 40 million clusters or 80 million 150bp paired-end reads on a NovaSeq 6000 instrument (Illumina) at the University of Colorado Cancer Center Genomics Shared Resource.

#### Data processing

Illumina adapters and the first 12 base pairs of each read were trimmed using BBDuk and reads <50bp post trimming were discarded. Reads were aligned and quantified using STAR (19) against the Ensembl human transcriptome (GRCh38.p12 genome (release 96)). Ensembl genes were mapped to HGNC gene symbols using HGNC and Ensembl BioMart. Gene counts were generated using the sum of counts for transcripts of the same gene. Lowly expressed genes were removed if mean raw count <1 or mean CPM (counts per million) <1 for the entire dataset. Reads were normalized to CPM using the edgeR R package (20). Differential expression was calculated using the voom function in the limma R package (21). In addition to two-group comparisons, single drug comparisons for all cell lines were generated with cell line as a covariate (**Table S9**).

#### Alignment and transcript quantification

STAR --runThreadN 12 --runMode genomeGenerate --sjdbGTFfile Homo_sapiens.GRCh38.96.gtf --genomeFastaFiles Homo_sapiens.GRCh38.dna_sm.primary_assembly.fa

STAR --readFilesIn Read1.fastq.gz Read2.fastq.gz --readFilesCommand zcat --runThreadN 6 --alignEndsProtrude 13 ConcordantPair --outFilterScoreMinOverLread 0.66 --outFilterMatchNminOverLread 0.66 --outSAMtype BAM SortedByCoordinate --quantMode GeneCounts

### Pathway analysis

Gene set enrichment analysis was performed using the full list of genes ranked by fold change for the indicated comparison and the fgsea R package (22) using gene sets from the Molecular Signatures Database (v7.0) (23). General plots were generated with the ggplot2 and ggpubr R packages (24). Heatmaps were generated with the ComplexHeatmap R package following z-score transformation (25).

### Proteomics

#### Sample preparation

All cell lines were grown for several passages in the absence of antibiotics, gemcitabine or cisplatin, then seeded at 100,000 – 200,000 cells per well and grown for 48 hours in IMDM + 10% FBS. Approximately 48 hours after seeding cells the supernatant was aspirated and cells were washed 3 times with cold phosphate-buffered saline (PBS). Cells were lysed in 100μl of 8M Urea, 50mM Tris-HCl, pH 8.0. Lysates were transferred to pre-chilled 1.5mL microcentrifuge tubes and centrifuged at 15000 RCF for 10 minutes to pellet. The supernatant was then transferred to a clean, pre-chilled tube and frozen. Lysate replicates were collected in triplicate from different passages. Cell pellets were lysed in 8M Urea supplemented with 0.1% Rapigest MS compatible detergent. DNA was sheared using probe sonication, and protein concentration was estimated by BCA (Pierce, Thermo Scientific). A total of 30μg protein per sample was aliquoted, and samples were diluted to <2M Urea concentration using 200mM ammonium bicarbonate while also undergoing reduction with DTT (10mM) and then alkylation with IAA (100mM). The pH of diluted protein lysates was verified as between 7-8, and samples were digested with sequencing grade Trypsin/Lys-C enzyme (Promega) in the presence of 10% Acetonitrile for 16 hours at 37°C. Samples were acidified adding formic acid to 1%, and speed vac dehydration was used to evaporate acetonitrile. Peptides were desalted on C18 tips (Nest group) and dried to completion. Prior to MS, peptides were resuspended in 0.1% Formic Acid solution at 0.5μg/μL concentration with 1:40 synthetic iRT reference peptides (Biognosys).

#### Data acquisition

Peptides were analyzed by liquid chromatography coupled with mass spectrometry in data independent acquisition (DIA) mode essentially as described previously (26). Briefly, 4μL of digested sample were injected directly unto a 200 cm micro pillar array column (uPAC, Pharmafluidics) and separated over 120 minutes reversed phase gradient at 1200 nL/min and 60°C. The gradient of aqueous 0.1% formic acid (A) and 0.1% formic acid in acetonitrile (B) was implemented as follows: 2% B from 0 to 5 min, ramp to 4% B at 5.2 minutes, linear ramp to 28% B at 95 minutes, and ramp to 46% B at 120 minutes. After each analytical run, the column was flushed at 1200 nL/min and 60°C by injection of 50% Methanol at 95% B for 25 minutes followed by a 10 minutes ramp down to 2% B and a 5 minute equilibration to 2% B. The eluting peptides were electro sprayed through a 30 μm bore stainless steel emitter (EvoSep) and analyzed on an Orbitrap Lumos using data independent acquisition (DIA) spanning the 400-1000 m/z range.

Each DIA scan isolated a 4 m/z window with no overlap between windows, accumulated the ion current for a maximum of 54 seconds to a maximum AGC of 5E5, activated the selected ions by HCD set at 30% normalized collision energy, and analyzed the fragments in the 200-2000m/z range using 30,000 resolution (m/z = 200). After analysis of the full m/z range (150 DIA scans) a precursor scan was acquired over the 400-1000 m/z range at 60,000 resolution.

#### Peptide library generation

To construct a comprehensive peptide ion library for the analysis of human BCa we combined several datasets, both internally generated and external publicly available data resources were utilized. First, we utilized a previously published (27) human bladder tumor proteomics experiment by downloading raw files from the online data repository (ProteomeXchange, PXD010260) and searching them through our internal pipeline for data dependent acquisition MS analysis (28) against the UniProt human reviewed canonical sequence database, downloaded July 2019, using internal peptides to perform retention time alignment (29). To this library, we appended a sample specific library generated from DIA-Umpire extraction of pseudo-spectra from one full set of replicates from the experimental bladder tumor cell lines. A final, combined consensus spectrast library containing all peptide identifications made between the internal and external dataset was compiled and decoy sequences were appended.

#### Data analysis

Peptide identification was performed as previously described in (28,29). Briefly, we extracted chromatograms and assigned peak groups using openSWATH (30) against the custom BCa peptide assay library described above. False discovery rate for peptide identification was assigned using PyProphet (31) and the TRIC (32) algorithm was used to perform feature-alignment across multiple runs of different samples to maximize data completeness and reduce peak identification errors. Target peptides with a false discovery rate (FDR) of identification <1% in at least one dataset file, and up to 5% across all dataset files were included in the final results. We used SWATH2stats to convert our data into the correct format for use with downstream software MSstats (33). Each individual data file was intensity normalized by dividing the raw fragment intensities by the total MS2 signal. MSstats (33) was used to convert fragment-level data into protein-level intensity estimates via the ‘quantData’ function, utilizing default parameters with the exception of data normalization, which was set to ‘FALSE’. For plotting purposes, protein intensities were VSN normalized, log-transformed, and replicate batch effects were removed using the removeBatchEffect function in the limma R package. The limma package was also used to calculate differential protein expression (21). Multiple hypothesis correction was performed using the Benjamin Hochberg method.

### Whole exome sequencing

#### Sample preparation

All cell lines were grown for several passages in the absence of antibiotics, gemcitabine, or cisplatin. Cell pellets were snap frozen from sub-confluent dishes for each of the 20 cell lines sequenced (5 cell lines, each with 4 derivatives: parental, Gem-resistant, Cis-resistant, GemCis-resistant). gDNA isolation was performed using the Puregene cell and tissue kit (Qiagen) with the addition of RNase A Solution (Qiagen) according to manufacturer’s instructions. gDNA was quantified using a Qubit 4.0, then sheared using a Covaris S220 Sonicator to 200bp. Libraries were constructed using the Sure Select All Exon v6 library kit (Agilent) following the XT library preparation workflow. Completed libraries were run on the 4200 Tape Station (Agilent) using D1000 screen tape. Libraries were quantitated using the Qubit, diluted to 4nM prior to verification of cluster efficiency using qPCR, then sequenced on the NovaSeq 6000 instrument (Illumina) (150bp, paired-end) at the University of Colorado Cancer Center Genomics Shared Resource. Mean insert size across all cell lines was 177.8 bp and mean coverage was 193.7X with > 96.8% at >30X. Individual call line quality control metrics are reported in **Table S11**.

#### Data processing

The analysis pipeline was developed using Nextflow. For the raw fastq files, Fastqc was used to assess overall quality. For computational efficiency, raw sequence reads were partitioned using BBMap (partition.sh) into 40 partitions. They then were aligned to the GRCh38 reference genome (including decoy sequences from the GATK resource bundle) using the BWA-MEM short read aligner (34), and merged back into single BAM files using Samtools. The resulting BAM files were de-duplicated using Samblaster (35), and sorted using Samtools. These duplicate-marked bams were further passed through the GATK Base Quality Score Recalibration in order to detect systematic errors made by the sequencing machine when it estimates the accuracy of base calls. The dbSNP (version 146) (36), the 1000 Genome Project Phase 1 (37), and the Mills and 1000G gold standard sets (38) were used as databases of known polymorphic sites to exclude regions around known polymorphisms from analysis. After alignment, Samtools (39), Qualimap (40), and Picard tools (41) were run to acquire various metrics to ensure there were no major anomalies in the aligned data.

#### Alignment

bwa mem -K 100000000 -R “read_group” -t 64 -M ref_fasta read_1 read_2

#### Marking duplicates

samtools sort -n -O SAM sample_bam | samblaster -M --ignoreUnmated

#### Base Quality Score Recalibration

gatk BaseRecalibrator -I sample_bam -O sample.recal.table -R ref_fasta --known-sites known_sites

#### Whole exome sequencing variant calling

We used Mutect2 from the GATK toolkit for SNVs and short indels (42). Mutect2 is designed to call somatic variants and makes no assumptions about the ploidy of samples. It was run in *tumor-only* mode to maximize the sensitivity albeit at the risk of high false positives. We used tumor-only mode to call variants for each cell line separately. Mutect2 workflow is a two steps process. In the first step, it operates in high sensitivity mode to generate intermediate callsets that are further subjected to filtering to generate the final variant calls. Annotation of variants was performed using Annovar (43) with the following databases: refGene, cytoBand, exac03, avsnp150, clinvar_20190305, gnomad211_exome, dbnsfp35c, cosmic90. Intergenic variants were removed along with variants that were identified at greater than 0.001% of the population according to ExAC or gnomAD, or had a depth < 20.

#### Mutect2 raw callset

gatk Mutect2 -R ref_fasta -I bam_tumor -tumor Id_tumor --germline-resource germline_resource

#### -O raw_vcf

Mutect2 filtering:

#### gatk FilterMutectCalls -V raw_vcf --stats raw_vcf_stats -R ref_fasta -O filtered_mutect2_vcf

Copy number calling using GATK Base quality score recalibrated bams were used as the input. The covered regions for the exome kit were converted into bins (defining the resolution of the analysis) for coverage collection. Read-counts, that form the basis of copy number variant detection, were collected for each bin. The read-counts then go through denoising, modelling segments, and calling the final copy ratios.

#### Preprocess intervals

gatk PreprocessIntervals --intervals intervals_bed_file --padding 0 --bin-length 0 -R ref_fasta --interval-merging-rule OVERLAPPING_ONLY -O preprocessed_intervals_list

#### Collect read counts

gatk CollectReadCounts -I sample_bam -L preprocessed_intervals} --interval-merging-rule OVERLAPPING_ONLY -O sample.counts.hdf5

#### Denoise read counts

gatk DenoiseReadCounts -I sample.counts.hdf5 --standardized-copy-ratios sample_std_copy_ratio --denoised-copy-ratios sample_denoised_copy_ratio

#### Model Segments

gatk ModelSegments --denoised-copy-ratios denoised_copy_ratio --output-prefix id_sample -O output_dir

#### Call copy ratio segments

gatk CallCopyRatioSegments -I sample.modelled_segments -O sampled.called.segments

#### Cell line authentication

Variant calls from the Mutect2 pipeline were filtered for each cell line to identify high confidence variants according to the filtering criteria above. These high confidence variants were then compared to the variants reported for all cell lines in the DepMap (https://depmap.org/portal/) for the Cancer Cell Line Encyclopedia (CCLE_mutations_hg38.csv, sample_info.csv) and COSMIC (CosmicCLP_MutantExport.tsv) as measured by the jaccard distance, the intersection of variants divided by the union of variants. Cells listed in CCLE or COSMIC were the rank ordered for each BCa cell line in this study according to the jaccard distance. Results are reported in **Table S12**.

### Cell Line Drug Treatments

Gemcitabine (Sigma) and cisplatin (Sigma) stocks were resuspended in 0.9% PBS solution. All stock solutions were stored protected from light and kept frozen until use. For cell culture dose response, cells were seeded in 96-well tissue culture plates with 500-2000 cells per well depending on the growth rate and duration of the experiment. Cells were seeded and allowed to attach overnight followed by replacing the media with fresh, pre-warmed media just prior to treatment. Drug dilutions were performed serially and using complete media (IMDM + 10% FBS) and the associated drug treatments. Growth inhibition was measured using confluence estimates over time on the IncuCyte ZOOM (Essen Bioscience) over varying amounts of time depending on each experiment. Dose response curves were generated with Prism v9.3.1 using a variable slope, four parameter nonlinear regression model. Comparison between treatment groups was done between IC50 values using the sum-of-squares F test. Details for timing and replicates for each experiment are included in their respective figure legends.

### Antibodies and Western Blotting

Whole cell lysates were prepared from cultured cells using RIPA lysis and extraction buffer (ThermoScientific). Lysates from xenograft tissues were prepared using tissue protein extraction reagent (T-PER) and glass tissue homogenizer. All lysates were prepared on ice and with the addition of Halt protease and phosphatase inhibitor cocktail and EDTA (ThermoFisher). Protein concentration of lysates were quantified with BCA protein assay (Pierce™, ThermoFisher). All lysates were prepared with 4X Licor Loading buffer with 50mM DTT added boiled for 10 minutes prior to gel loading. All western blots were run using PROTEAN TGX precast 4-15% or 4-20% gradient gels (Bio-Rad) and transferred to either 0.2μm or 0.44μm nitrocellulose membranes. Transfer was done for 1.5-2hrs in cold TrisGlycine buffer (Bio-Rad) with 20% methanol prior to blocking for 1hr at room temperature in 5% BSA in TBS-T. Primary antibodies were diluted and incubated overnight at 4°C on a rocker. Membranes were washed 3 or 4 times in fresh TBS-T prior a 1-hour room temperature incubation in an appropriate secondary antibody. Membranes were washed 3-4 times in TBS-T, developed with enhanced SuperSignal West Pico Plus or SuperSignal West Fempto (ThermoFisher), and imaged using Li-Cor Odyssey^®^ Fc instrument. Densitometry was performed using LiCor Image Studio™ software. Statistical comparisons using densitometry measurements were done using a one-way ANOVA with Tukey post hoc to control for the experiment-wise error rate.

Western Blot Primary Antibodies:

| Target | Vendor | Catalog number | Dilution | Type |
| --- | --- | --- | --- | --- |
| GAPDH | Cell Signaling | 5174S | 1:1000 | Rabbit polyclonal |
| H2AX | Thermofisher | MA1-2022 | 1:1000 | Mouse monoclonal |
| NPEPPS | Thermofisher | PA5-83788 | 1:1000 | Rabbit polyclonal |

Western Blot Secondary Antibodies:
| Target | Vendor | Catalog number | Dilution | Type |
| --- | --- | --- | --- | --- |
| Goat-anti-Rabbit(HRP) | Abcam | ab97057 | 1:20,000 | Goat polyclonal |
| Rabbit-anti-Mouse(HRP) | Sigma | A9044-2ML | 1:10,000 | Rabbit polyclonal |

### shRNA-mediated knockdown experiments

Lentiviral production and transduction were carried out by the University of Colorado Cancer Center Functional Genomics Shared Resources. Plasmids from The RNAi Consortium (TRC) collection (TRC construct numbers TRCN0000073838, TRCN0000073839 and TRCN0000073840) were used for targeting NPEPPS were selected based on predicted knockdown efficiency; non-targeting controls used were SHC002 and SHC016. 2μg of target shRNA construct and 2μg of 3:1 ratio of psPAX2 (Addgene) and pMD2.G (Addgene) were transfected into HEK293FT cells using 2 μg of Polyethylenimine (Polysciences). Lentiviral particle containing media was filtered using 0.45μm cellulose acetate syringe filter and used for transduction. Puromycin selection was performed at doses used for CRISPR library screening or in some cases, cells were re-selected with higher doses of puromycin (10μg/mL), in order to ensure the complete elimination of non-transduced cells. Selected cells were frozen at early passage and early passage cells were used for all experiments.

### Intracellular cisplatin measurements using CyTOF

Cell lines were cultured for several passages in IMDM + 10% FBS. Prior to the experiment, cells were cultured in IMDM10 to be 50-80% confluence overnight and then treated the next day with 10μM cisplatin or PBS and then dissociated after 4 hours of treatment. For dissociation, cells were washed twice with room temperature PBS and then incubated with PBS + 0.05% Trypsin-EDTA for 10-15 minutes. Cells were neutralized with IMDM10 and then fully dissociated into single-cell suspension by gentle pipetting. After dissociation, cells were counted by Trypan blue staining and then placed in separate tubes at 3 x 10^5^ cells. Individual samples were then fixed, permeabilized, and labeled using unique barcodes using the Cell-ID 20-plex Pd Barcoding kit (Fluidigm) according to the manufacturer’s protocol. Barcoded samples were pooled across cell line conditions and cisplatin concentration, incubated with Cell-ID Intercalator-Ir, mixed with equilibration beads, and acquired on a Helios mass cytometer (Fluidigm). Post-acquisition data were normalized to equilibration beads and debarcoded, using the bead-normalization and single-cell-debarcoder packages from the Nolan Laboratory GitHub page (https://github.com/nolanlab). Relative cisplatin intensity (defined by ^195^Platinum isotopic mass intensity) was analyzed among nucleated ^191^Iridium+ ^193^Iridium+ events defined by Boolean gating within FlowJo 10.7.1.

### Whole Genome CRISPR Screening

#### Plasmid library expansion and quality control

Whole genome CRISPR Screening was performed using the Human CRISPR Knockout Pooled Library (Brunello) - 1 vector system (Addgene and a gift from John Doench to the Functional Genomics Facility at the University of Colorado Anschutz Medical Campus)(44). Two distinct plasmid expansions were performed. And the library distribution was assessed using next generation sequencing to determine the impact on overall library was modest following re-expansion. Library width was calculated as previously described(45,46) by dividing the 10^th^ percentile of the library distribution by the 90^th^ percentile using the log2 average expression of all sgRNAs in the library and found to be 6.7 and 7.13 for batch 1 and 2 respectively. All quality control metrics for each sample are reported in **Table S13**. Different screening parameters were used based on the cell line screened these are summarized in **Table S4.**

#### Lentivirus Production and Titration

For the two plasmid batches, two distinct protocols for lentivirus production were utilized. The first batch was generated by using Polyethylenimine, linear (PEI; Polysciences) and was used for the T24-GemCis and TCCSUP-GemCis screens. The second used lipofectamine 3000 and was applied for the 253J-GemCis, KU1919-GemCis, and 5637-GemCis screens. For the first batch, 293FT cells were seeded at a density of 36,800 cells/cm^2^ into a 4-layer CELLdisc (Greiner) using DMEM + 10% FBS along with antibiotic and antimycotic solution. Transfection mix consisting 47.6μg pMD2G (Addgene), 95.2μg of psPAX2 (Addgene), and 190.5μg of Brunello Whole genome knockout library (Addgene) was mixed with 448μl PEI (1 mg/mL) and 3mL OptiMEM, vortexed for 30 seconds and allowed to incubate at room temperature for 20 minutes. Fresh media containing transfection mix were added to the CELLdisc using up to 270mL of media. The next day media was changed for 280mL fresh media followed by a 48-hour incubation. After this 48-hour incubation the viral supernatant was harvested and filtered through a cellulose acetate filter system (ThermoScientific) and frozen at -80°C.

The first method had low functional virus titer, so we implemented a different virus production method for subsequent screens. In the second batch of virus production, we utilized lipofectamine 3000 instead of PEI, eliminated use of multilayer flasks and centrifuged to remove debris as opposed to filtering. Briefly, 293FT cells were plated in T225 flasks to be 80% confluent after 24hrs. 2hrs before transfection, media was changed and 40mL of fresh media was used per T225 flask. The lipofectamine 3000 protocol was followed according to manufacturer’s instructions and scaled based on the volume of virus being prepared. For each T225 flask 2mLOptiMEM was mixed with 40μg Brunello whole genome library plasmid, 30μg of psPAX2 and 20μg of pMD2.G and 180μl of P3000. This mix was added to a tube containing 2mL OptiMEM and 128μl Lipofectamine 3000, which was scaled according to the number of T225 flasks being prepared. Transfection mix was mixed thoroughly by pipetting up and down slowly, and allowed to incubate at room temperature for 15 minutes. Transfection mix was then added dropwise to the plates of 293FT cells with gentle swirling and incubated overnight (∼16hr). The following morning, the media was changed and 60mL of fresh media was added to each T225 flask. This was allowed to incubate overnight and replaced the following morning.

This first lentiviral supernatant was stored at 4°C to be pooled with a subsequent 48 hour collection. Upon collection, viral supernatants had 1M HEPES added at 1%. Following the second virus collection, supernatants were pooled and centrifuged at 1250rpm for 5 minutes to pellet debris. Lentivirus was stored in polypropylene tubes as polystyrene is known to bind lentivirus, and all tubes were flash frozen in liquid nitrogen and stored at −80°C. Despite the changes to the lentiviral production protocols, functional lentiviral titers were not improved using these changes to the methodology, but feel it is worth noting these changes in protocol to account for any possible variability associated with this change.

Lentivirus was titered functionally based on protocols adapted from the Broad Institute’s Genetic Perturbation Platform’s public web portal (https://portals.broadinstitute.org/gpp/public/).

#### Screening Parameter Optimization

All screening parameters for each cell line including cell polybrene and puromycin sensitivity, screening coverage, technical and biological replicates performed, and gemcitabine and cisplatin treatment concentrations are reported in **Table S4.**

#### DNA Isolation

Cell pellets of 2e7 were snap frozen in liquid nitrogen in 1.5mL tubes and stored at −80 prior to extraction. When possible, at least 8e7 cell were used for 4 separate genomic DNA isolation which were pooled to account for any variation with pellet size. DNA isolation was performed using the Puregene cell and tissue kit (Qiagen) with the addition of RNase A Solution (Qiagen) according to the manufacturer’s instructions. DNA concentration was measured in quadruplicate using either a nanodrop spectrophotometer (Thermo), Qubit® dsDNA assay (Life Technologies), and the average DNA content per cell was determined.

#### Library preparation

The minimum number of cell equivalents of gDNA to maintain equal coverage was used for library preparation. In all screens, the minimum coverage based on cell number was multiplied by the average gDNA content per cell for each individual cell line to determine the minimum number for 10μg PCR reactions needed to maintain coverage. A minimum coverage of 500-fold per sgRNA in the library was targeted for each independent sample or replicate but this was increased in some cases where screening was carried out with greater depth (see **Table S4** for coverage and replicate information).

Library preparation was performed using primers sequences designed by the Broad Institute’s Genetic Perturbation Platform (https://portals.broadinstitute.org/gpp/public/) and utilized a pool of eight P5 primers with to introduce a stagger in reads associated with each library and sample specific P7 primer that contained a unique sample index sequence for each timepoint, replicate, or treatment condition to be sequenced in the same pool (**Table S14**). All library preparation primers were resuspended at 100μM.

Each library preparation PCR reaction contained the following components: 1μl Herculase II Fusion Enzyme (Agilent), 2.5μl Deoxynucleotide (dNTP) Solution Mix (New England Biolabs), 0.5μl P5 primer pool, 0.5μl P7 index primer, 20μl 5X Reaction Buffer (Agilent), 10μg of gDNA and nuclease-free water to bring the total reaction volume to 100μl. Samples underwent 23 cycles of thermal cycling followed by a quality assessment by electrophoresis on 2% agarose gel to ensure consistent library amplification across multiple wells and samples for each plate.

Each unique library had 10μl pooled from all PCR reactions performed on that unique sample and mixed thoroughly. 50-100μl of the pooled library preparation reactions was used to perform magnetic bead-based purification and elimination of any residual free primer using a 0.8X ratio SPRIselect beads (Beckman Coulter) according to the manufacturer’s instructions. Libraries were then assessed for appropriate amplicon size and complete elimination of free primer peaks using the High Sensitivity ScreenTape Assay on the Tape Station2200 (Agilent) and quantified using the qPCR-based quantification in order to ensure only NGS-compatible amplicon was quantified using the Library Quant ROX Low Kit (Kapa Biosystems) on a QuantStudio™ 6 Realtime PCR System (ThermoFisher). Following qPCR quantification, all libraries were normalized to a standard concentration (typically 20-40nM) depending on the lowest concentration library to be pooled, and then requantified by qPCR to ensure all samples were within ∼10-20% of the pool mean target concentration. After confirming accurate library quantification and normalization, samples were pooled at an equimolar ratio and submitted for sequencing. Libraries were sequenced on the NovaSeq 6000 instrument (Illumina) (150bp, paired-end) at the University of Colorado Cancer Center Genomics Shared Resource.

#### CRISPR screening bioinformatic pipeline and analysis

sgRNA counts were extracted directly from R1 raw sequence reads using a custom perl script that uses regular expression string matching to exactly match sgRNA sequence flanked by 10 bases of vector sequence. The vector sequence was allowed to have one error before and after the sgRNA sequence. sgRNAs were tabulated for each sample based on the sgRNA sequence (**Table S15**). The sgRNA IDs of the Brunello library were updated to current HGNC gene names using the Total Approved Symbols download from HGNC, accessed on 9/1/2020 (https://www.genenames.org/download/statistics-and-files/). Transcript IDs were matched when possible and when matches were not found, past symbols and aliases were updated to current names. Finally, 5 sgRNAs with missing updated gene names were manually curated using literature searches. Library distribution was calculated using the caRpools R package (47). The DESeq2 R package (48) was used to calculate differential abundance of genes (**Table S5**). Gene counts were generated using the sum of counts for sgRNAs of the same gene. Synthetic lethality compared GemCis day 19 and GemCis day 25 vs. PBS day 19 and PBS day 25 with the day as a covariate. In the comparison integrating all cell lines, cell line was additionally modeled as a covariate. Gene essentiality was calculated by comparing PBS day 25 to PBS day 0 and in the integrated all cell lines comparison; cell line was modeled as a covariate. Common synthetic lethal genes were defined as being statistically significantly differentially lost (FDR < 0.05 and Log2 FC < 0) in each of the 5 cell lines. Gene set enrichment analysis (GSEA) was performed using the fgsea R package run with 10000 permutations (22) with the KEGG and Reactome gene sets from MSigDB (23). Heatmaps were generated with the ComplexHeatmap R package following z-score transformation (25). Other plots were generated using the ggplot2 R package.

### Xenograft experiment

Six-week-old, female NU/J mice (Jackson Labs) were allowed to acclimate for at least one week prior to initiating any experiments. Mice had free access to food and water in pathogen-free housing and cared for in accordance NIH guidelines and all experiments were performed under protocols approved by the University of Colorado Denver Institutional Animal Care and Use Committee (IACUC).

For KU1919-GC xenografts, cells that had been stably transduced with non-targeting control (shCtrl1, SHC002) and NPEPPS (shN39, TRCN0000073839) shRNA constructs. Mice were divided into groups of 22 and 23 for the non-targeting control and NPEPPS shRNA constructs respectively. Mice were injected with 4e6 cells in phenol red- and serum-free RPMI mixed with equal volume Matrigel Matrix (Corning) to total 100μl volume. Tumors were allowed to engraft for 9 days following injection and mice were randomized based on tumor volume within each shRNA condition into groups of 11 or 12 to be treated with combination gemcitabine plus cisplatin or DPBS. Treatment was initiated 13 days post-inoculation with dosing adjusted based on individual mouse weight.

Cisplatin (Sigma) and gemcitabine hydrochloride (BOC Sciences) were both resuspended in 0.9% PBS and stored protected from light at −80°C as individual aliquots. Prior to treatment fresh aliquots of gemcitabine and cisplatin were thawed and diluted to their final concentration with 1X DPBS (Gibco). Mice were treated three times weekly on a Monday, Wednesday and Friday schedule for four weeks total. All mice in the gemcitabine plus cisplatin-treated groups were given 50mg/kg gemcitabine and 2mg/kg cisplatin that were mixed and administered as a single intraperitoneal injection, while control mice were administered an equivalent volume of DPBS.

Mouse health was monitored daily and all tumor volume measurements and weights were measured 3x weekly schedule. Tumor volume was calculated using the formula (*L* x *W*^2^)/2, for which *L is* the length of the long axis and *W* is the width of the axis perpendicular to the long axis measurement. All measurements were performed using digital calipers. Animals were humanely euthanized with CO_2_ followed by cervical dislocation when tumors reached a predetermined endpoint of 2cm^3^ or when weight loss exceeded 15% body weight. Mice that were removed from the study due to weight loss were censored in the survival analyses.

### Linear mixed-effects model of tumor growth

Linear mixed-effects models were used to model longitudinal observations of xenograft tumor growth volumes normalized by their corresponding baseline volume. Mixed-effects models from the R-package *lme4* (49) and Satterthwaite’s approximation for degrees of freedom for the fixed effects from *lmerTest* (50) were used for model fitting and inspection in the R statistical software (4.0.3). Volume changes compared to baseline were log_2_-transformed. The final model was structured as:

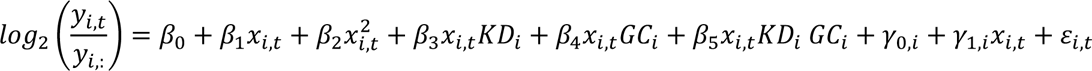

where *β* is the fixed effects capturing population-level trends, *γ* is the normally distributed random effects capturing individual-level variation, *ε* is the i.i.d. normally distributed residual term, *i* is the unique individual identifier, *t* notes the time points, *x_i,t_* ∈ {2, 4, 5, 7, 9, 11, 14, 16, 18, 21, 23, 25, 28} depicted days since initiating interventions, *y*_”,:_ is tumor volume at baseline prior to treatments upon randomization, and *y*_”,$_ were the observed tumor volumes over the treatment period measured in mm^3^. The model was fit using Restricted Maximum Likelihood and built iteratively until the underlying model assumptions and model convergence criteria were met. To this end, a quadratic growth term (*β*_2_) was added on top of the linear growth term (*β*_1_) and intercept (*β*_0_), allowing slightly non-linear relative growth patterns to be captured by the otherwise linear model. Binary indicators *KD_i_* ∈ {0,1} and *GC_i_* ∈ {0,1} were used to model knockdown of NPEPPS, GemCis treatment, or the combination. The corresponding model terms were captured in *β*_3_, *β*_4_ and *β*_5_, respectively. Finally, the model allows for individual-specific random effects for intercept (*γ*_0,i,_) and linear growth slope (*γ*_1,i_). Shapiro-Wilk test was used to examine the underlying normality assumption for *γ*_0,i_ and *γ*_1,i,_ with *p*=0.1373 and *p*=8901, respectively, indicating that these random effects followed underlying assumptions of normality. After inspection of the residual plots (**Figure S7B**), this final model was deemed suitable for population-level statistical inference via the fixed effects. This population-level model fits are visualized in **Figure S7A**. These population-level estimates are as follows:

| Fixed effect | Estimate | Std. error | df | t | p-val |
| --- | --- | --- | --- | --- | --- |
| $\beta_0$ (intercept) | 0.05054 | 0.08422 | 54.28 | 0.600 | 0.55091 |
| $\beta_1$ (linear slope) | 0.1236 | 0.01493 | 65.52 | 8.276 | 8.92e-12 *** |
| $\beta_2$ (quadratic slope) | 0.00308 | 0.0002242 | 389 | 13.740 | < 2e-16 *** |
| $\beta_3$ (knockdown) | -0.0605 | 0.01821 | 44.97 | -3.322 | 0.00178 ** |
| $\beta_4$ (GC) | -0.1063 | 0.01821 | 44.97 | -5.837 | 5.49e-07 *** |
| $\beta_5$ (knockdown + GC) | -0.1233 | 0.01791 | 45.28 | -6.884 | 1.47e-08 *** |

### Patient tumor-derived organoids (PDOs)

#### Culture of the PDOs

Human bladder tissue was obtained from the Erasmus MC Bladder Cancer Center, Rotterdam, the Netherlands and the Amphia Ziekenhuis, Breda, the Netherlands. Bladder tumor-derived organoids from biopsies obtained through TURBT or cystectomy were isolated and cultured using methods developed by Mullenders et al. (51) with modifications (Akbarzadeh/Scholtes et al. in prep). Briefly, bladder tissues were washed with Advanced DMEM/F12 (Gibco) supplemented with 10mM HEPES (Gibco), 1% GlutaMax (Gibco) and 100 μg/ml primocin (InvivoGen), henceforth Ad+++. Tissue was minced and incubated at 37°C with the digestion solution (collagenase 2.5mg/ml in EBSS) and isolated cells were passed through 70μM strainer (Falcon), washed with Ad+++ and seeded in 50 µl drops of BME (R&D system) containing 10000-15000 cells in 24 well suspension plates (Greiner). Bladder tumor organoids were cultured in a culture medium containing Ad+++ supplemented with 1 × B-27 (Gibco), 1.25 mM N-acetylcysteine (Sigma), 10 mM nicotinamide, 20μM TGF*β* receptor inhibitor A83-01, 100ng/ml recombinant human FGF10 (Peprotech), 25 ng/ml recombinant human FGF7 (Peprotech), 12.5 ng/ml recombinant human FGF2 (Peprotech), 10μM Y27632 Rho Kinase (ROCK) Inhibitor (Sigma) and conditioned media for recombinant Rspondin (2.5% v/v), and Wnt3A (2.5% v/v). The medium was changed every three days. Organoids were passaged at a 1:3 to 1:6 ratio every 7 days using cell dissociation solution-non enzymatic (Sigma) and plated in fresh BME matrix droplets.

#### Lentiviral transduction of bladder cancer PDOs

Lentiviral vectors containing the desired shRNA sequences were amplified from bacterial glycerol stocks obtained in house from the Erasmus Center for Biomics and part of the MISSION® shRNA library. The same shRNA constructs used for bladder cancer cell lines, and described above, were used for PDOs. For overexpression, pLenti-C-Myc-DDK-P2A-Puro Lentiviral Gene Expression Vectors were acquired from Origene (NPEPPS RC209037L3, empty vector PS100092). For lentiviral particle generation, in total, 5e6 HEK293T cells were plated in a 10-cm dish and transfected with 12.5 μg of plasmids mix. In total, 4.5 μg of pCMVΔR8.9 (envelope), 2 μg of pCMV-VSV-G (packaging) and 6 μg of shRNA vector were mixed in 500 μl serum-free Opti-MEM and combined with 500 μl Opti-MEM containing 125 μl of 10 mM polyethyleneimine (PEI, Sigma). The resulting 1 ml mixture was added to HEK293T cells after 15 min incubation at room temperature. The transfection medium was removed after 12 h and replaced with a fresh RPMI medium. Virus-containing medium was harvested and replaced with fresh medium at 48, and 72 h post-transfection. After each harvest, the collected medium was filtered through a cellulose acetate membrane (0.45 μm pore), viral particles were concentrated by ultracentrifugation at 20.000 rounds per minute for 1 hour at 4 °C with pellets re-suspended and pooled in a final volume of 2mL adv. DMEM for each condition. Medium containing viral particles was used either directly or was stored at −80 °C. For lentiviral transduction of bladder cancer organoid cells, organoids were harvested and dissociated to single cells using cell dissociation solution. For each condition (shRNA control / shRNA NPEPPS / NPEPPS-OE / empty vector), 1 × 106 cells were collected and gently re-suspended in 1mL of concentrated lentivirus. Cell-virus mixture was then divided over two wells of a pre-warmed 24-well plate and subsequently sealed with parafilm. Plates were spinoculated at 600 x g at 25 °C for 1 hour. Parafilm was removed immediately after spinoculation, and plates were incubated for 6-8 hours at 37 °C 5% CO2. Transduced cells were washed 1 time with 10mL advanced DMEM spinning down at 200 x g for 10 minutes and seeded in domes at a density of 150 cells per μL BME. Once matured, organoids were treated with 2 µg/ml puromycin for 72 hours. Puromycin-selected organoids were then passaged and given 6 days of recovery time prior to experimental procedures.

#### Ex vivo drug testing

Organoids were collected 4-7 days after passaging and dissociated to single cells using a cell dissociation solution, assisted by mechanical dissociation. All assays were performed in 96 well suspension plates (Greiner bio-one; 655 185), seeding 10,000 cells per well in 100 µl bladder organoid medium containing 15% BME. Drugs were added when mature organoids formed after two-three days, and drug treatment was performed in triplicate. For *ex vivo* cisplatin response: cisplatin (Sigma; PHR1624; reconstituted in NaCl) was added in six doses in the range of 0.1 to 40 µM. Caspase-3 and −7 activity was measured following three days of treatment (Caspase-Glo #G8093, Promega), while cell viability was obtained after six days of treatment (CellTiter-Glo 3D #G9681). Plates were read on a SpectraMax I3 plate reader. Viability data was normalized using organoid wells treated with vehicle control (1.2% PBS) (Medchemexpress; HY-14807; reconstituted in DMSO). Organoids were harvested after six days of treatment and dissociated into single cells using a cell dissociation solution, assisted by mechanical dissociation. Three-quarters of the cells were used to obtain cell viability as previously described (treatment), while one-quarter of the cells were reseeded in 200µL BOM + 15% BME. Following reseeding, organoids were allowed to regrow for another six days after which cell viability (reseeding) was again obtained. Cell viability was normalized using organoids treated with vehicle control (0.02% DMSO, 1.2% PBS).

#### SNaPshot mutation analysis

Tumor and matched organoid DNA was isolated using with the QIAmp DNA Mini-Kit (Qiagen) according to the manufacturer’s protocol. Presence of hotspot mutations in the *TERT* promoter sequence chr5:1,295,228C>T, chr5:1,295,248G>A and chr5:1,295,250C>T [GRCh37/hg19]), *FGFR3* (R248Q/E, S249C, G372C, Y375C, A393E, K652E/M) and *PIK3CA* (E542K, E545G/K and H1047R) were assessed on tumor and organoid DNA by SNaPshot mutation analysis with the same methods as previously described (52–54).

#### Organoid phenotyping and tumor histology

Tissue processing and H&E staining were performed using standard procedures. For hematoxylin-eosin (H&E) staining of organoids, wells of BME-embedded organoids were fixated with 4% formalin (Sigma) and 0.15% glutaraldehyde (produced in-house) at room temperature for 2 hours. Fixated BME and organoids were washed with PBS and engulfed in 2.5% Low-Melting Agarose (Sigma) prior to paraffin embedding. H&E staining was performed on 4μM paraffin sections of both tumor and organoid tissue. Stained whole-slides, as well as prior 3D organoid cultures, were imaged by bright-field microscopy (Olympus IX70).

#### Data availability

The mass spectrometry proteomics data have been deposited to the ProteomeXchange Consortium via the PRIDE (55) partner repository with the dataset identifier PXD024742. The whole exome sequencing data have been deposited in the BioProject database with project identifier PRJNA714778. The RNA sequencing data have been deposited in the GEO database with dataset identifier (GSE171537). The CRISPR screen sequencing data have been deposited in the GEO database with dataset identifier (GSE179799). The copy number data have been deposited in the ArrayExpress database with identified (E-MTAB-10353). All data are made available through a custom build R Shiny app at: https://bioinformatics.cuanschutz.edu/BLCA_GC_Omics/

#### Reviewer Login Information

##### PRIDE

Username: Password: n70IwGNc

##### SRA

https://dataview.ncbi.nlm.nih.gov/object/PRJNA714778?reviewer=50g24ej0558qj2dpr9v6pekgvl

##### GEO (RNAseq)

https://www.ncbi.nlm.nih.gov/geo/query/acc.cgi?acc=GSE171537 token: avkdyoiixnettod

##### CRISPR Screen

URL: https://www.ncbi.nlm.nih.gov/geo/query/acc.cgi?acc=GSE179799 Reviewer token: elafaomerfarvsr

##### ArrayExpress

Username: Reviewer_E-MTAB-10353

Password: u3MpsspP

## RESULTS

We acquired the five human BCa cell lines, KU1919, 5637, T24, TCCSUP, and 253J from the Resistant Cancer Cell Line (RCCL) collection (56,57). For each, we obtained the parental lines (-Par) and derivatives made resistant through dose escalation to cisplatin (-Cis), gemcitabine (-Gem), and the combination of gemcitabine plus cisplatin (-GemCis) (**Figure 1A, Table S1**). We confirmed resistance to the associated drugs for all resistant derivatives in comparison to the parental lines and found them to be consistent with those reported by the RCCL (**Figure S1**) (56,57). These cells represent features and alterations in putative BCa drivers as reported in TCGA (58) and variants reported in ClinVar (59) (**Tables 1**, **S2, S3**).

### Genome-wide CRISPR screens identify 46 common synthetic lethal genes

To study the connection between drug resistance and gene expression, we performed whole-genome loss-of-function screens in each of the five GemCis-resistant cell line derivatives. After transduction of the Brunello CRISPR-Cas9 knockout library (44), we passaged the cells for 10 days to clear essential genes, then split them into phosphate-buffered saline (PBS) or gemcitabine plus cisplatin treatment groups (**Figure 1A**). Each screen was performed at a drug concentration that allowed the GemCis-resistant cells to grow unrestricted, but significantly inhibited the growth of the parental lines (**Table S1**). Screening parameters for each cell line are reported in **Table S4**. We counted sgRNAs 9 and 15 days after the start of treatment (19 and 25 days after transduction). As expected, similar experimental conditions clustered together when correlations were measured across treatment conditions and cell lines (**Figure S2**).

We defined genes as “synthetic lethal” with gemcitabine plus cisplatin treatment as those for which the combined cognate sgRNA counts were significantly lower (moderated t-test, FDR < 0.05) in the gemcitabine plus cisplatin-treated arm compared to the PBS arm when including both days 19 and 25 in the statistical model (**Table S5**). We identified 235 synthetic lethal genes that were statistically significant in KU1919-GemCis, 888 for T24-GemCis, 2099 for TCCSUP-GemCis, 2369 for 253J-GemCis, and 511 for 5637-GemCis. Next, we performed gene set enrichment analysis (60) on the full ranked list of genes according to their synthetic lethality. For this analysis, we created one ranked gene list by including each of the five cell types in the statistical model directly. As expected, and as a validation of the screen itself, we found that the top-ranked pathways were dominated by processes such as DNA repair, Fanconi Anemia, nucleotide excision repair, double-stranded break repair, base-excision repair, and DNA damage bypass mechanisms (**Figure 1B, Table S6**). These results are consistent with the known roles of DNA damage detection and repair in cisplatin resistance (61,62). In addition, the overall ranking of genes from our CRISPR screen showed enrichment for manually curated genes associated with platinum resistance in cancer (63) (**Table S5**).

From these results, we identified 46 genes that were commonly synthetic lethal across all five cell lines (**Figure 1C, S3)**. Consistent with the overall findings (**Figure 1B**), 41 of the 46 common synthetic lethal genes fell into one or more putative DNA damage response and repair pathways, including homologous recombination, double-stranded break repair, nuclear excision repair, and Fanconi anemia (**Figure S3B**, **Table S7**). These genes showed a range of growth patterns over the full length of the CRISPR screen. As illustrated in **Figure 1D**, some genes showed patterns of increased cell growth in PBS treatment, then reduced growth in gemcitabine plus cisplatin treatment. Other genes had very little impact on cell growth in PBS treatment, but then reduced growth when treated with gemcitabine plus cisplatin. Finally, some genes reduced cell growth in PBS treatment and further reduced growth with gemcitabine plus cisplatin treatment. Overall, these results provide a robust list of known and novel genes involved in chemotherapy resistance that can potentially be targeted therapeutically to improve treatment response.

**Table 1.** Clinicopathologic characteristics and genetic drivers for five cell lines.

| Feature | KU1919 | T24 | TCCSUP | 5637 | 253J |
| --- | --- | --- | --- | --- | --- |
| **Sex** | Male | Female | Female | Male | Male |
| **Stage** | T3 | Ta | N/A | N/A | T4 |
| **Grade** | G3 | G3 | G4 | G2 | G4 |
| **Base47 Subtype** | N/A | Basal | Basal | Luminal | Basal |
| **TP53** |  | Y126X | E349X |  |  |
| **HRAS** |  | G12V |  |  |  |
| **NRAS** | Q61R |  |  |  |  |
| **PIK3CA** |  |  | E545K |  | E545G |
| **TERT** |  |  |  |  |  |
| **ARID1A** | Y1052X |  |  |  |  |
| **KMT2D** | T2441Pfs*44 |  |  | Q2813X |  |
| **KDM6A** | Q915X |  |  |  |  |
| **FAT1** |  | S2682X | D1536N |  |  |
| **KMT2C** |  | R4225X;<br>A3559T |  |  |  |
| **ERBB2** |  |  |  | S310F |  |
| **ERBB3** |  | E1219K |  |  |  |
| **EP300** |  | C1201Y |  |  |  |
| **FBXW7** |  |  | S66X |  |  |
| **ASXL2** |  |  | E330Q |  |  |
| **ATM** |  |  |  | H1876Q |  |
| **AKT1** | E17K |  |  |  |  |
| **RYR2** |  | R2401H |  |  |  |
| **NFE2L2** |  |  |  |  | G81S |
| **RB1** |  |  | LOSS | Y325X |  |
| **E2F3** |  |  | AMP | AMP |  |
| **PPARG** |  |  |  | AMP |  |
| **CCND1** | AMP |  |  |  |  |
| **CDKN2A** | LOSS |  |  |  | LOSS |

### Complementary multi-omic profiling prioritizes NPEPPS as the top driver of treatment resistance

Pre-treatment multi-omic profiling has been shown to reveal the biological impact of synthetic lethal hits (64). Accordingly, we performed RNA sequencing and mass spectrometry-based proteomic profiling on cell lysates of all cell lines grown in drug-free media (**Figure 1A**). We leveraged transcriptome and proteome profiles from parental to matched drug-resistant derivative lines (-Gem, -Cis, and -GemCis) to prioritize the 46 common synthetic lethal genes that drive resistance in the context of gemcitabine plus cisplatin treatment resistance.

Taking this approach, the transcriptomics data revealed 1557 significantly upregulated genes across the Gem-resistant lines, 1897 in the Cis-resistant lines, and 1530 in the GemCis-resistant lines (moderated t-test, FDR < 0.05; **Table S9**). The proteomics data revealed 9 significantly upregulated proteins across the Gem-resistant cell lines, 1 in the Cis-resistant cell lines, and 10 in the GemCis-resistant cell lines (moderated t-test, FDR < 0.25; **Table S10**). Given the lower number of significant proteins and the relevance of transcript expression in predicting genetic dependency (64), we first investigated the overlap between the CRISPR screen results and the transcriptomes from each of the resistant cell line derivatives compared to the parental cells. Few genes were significantly and consistently upregulated across the resistant derivatives in the list of 46 commonly synthetic lethal genes (**Figure 2A**), but the most significantly and consistently upregulated genes were involved in DNA damage response and repair mechanisms, including ERCC6, XPA, REV1, POLH, ERRC8, PRIMPOL, NBN, and members of the Fanconi Anemia pathway. We identified Puromycin-Sensitive Aminopeptidase, NPEPPS, to be the most consistently upregulated gene across the resistant derivatives at the RNA level (**Figure 2A,B**). Similarly, we found NPEPPS to be consistently and significantly upregulated at the protein level (**Figure 2C**). NPEPPS was also a top synthetic lethal hit (**Figure 2D, Table S5**). Consistent with the proteomics results, immunoblotting for NPEPPS revealed that it was upregulated in the Cis-resistant and GemCis-resistant lines, with the Gem-resistant lines showing variable upregulation (**Figure 2E**).

**Figure 2.**
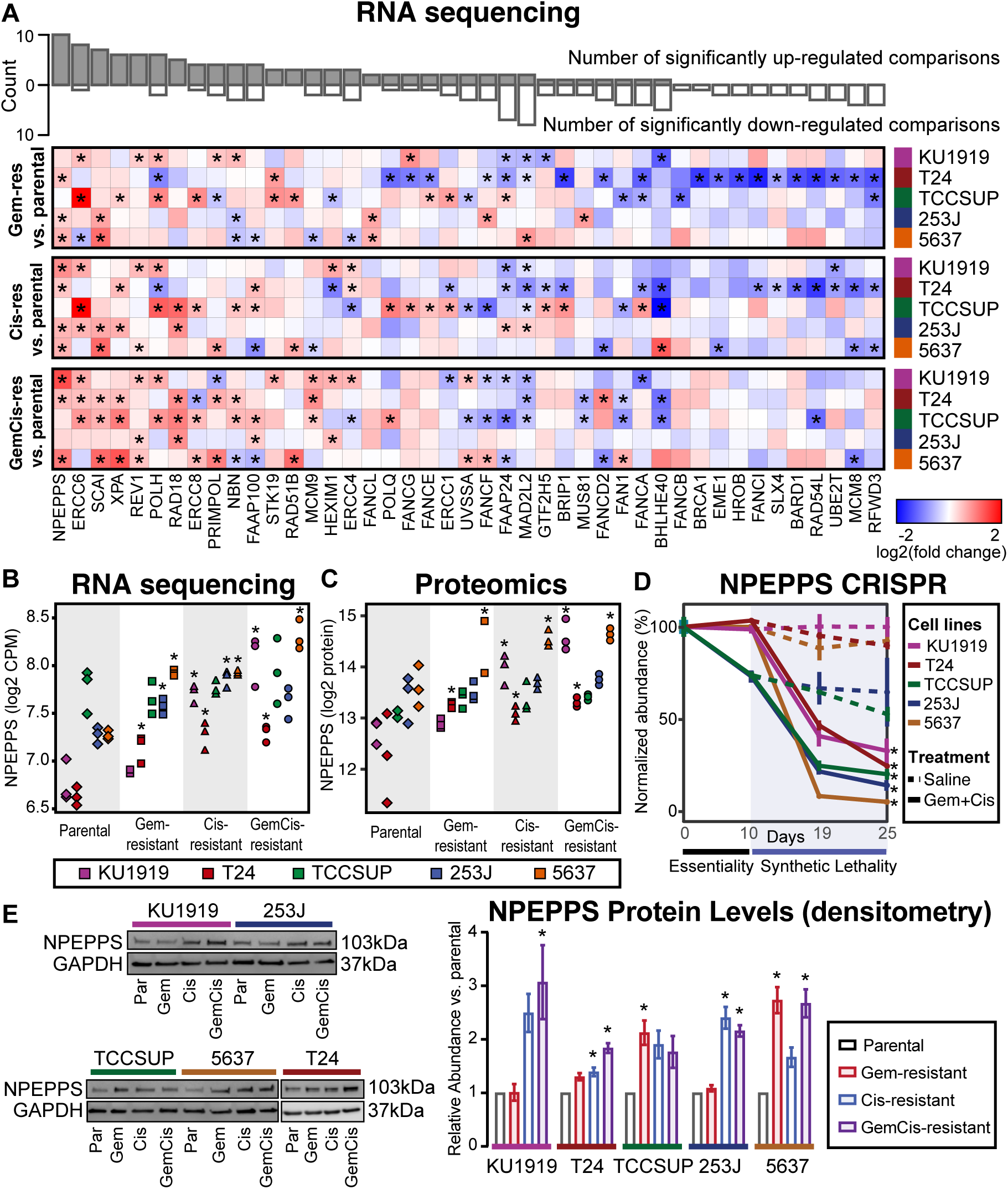
NPEPPS is identified as a commonly upregulated and synthetic lethal hit. (**A**) Differential gene expression of 43 common synthetic lethal genes as measured by RNAseq across all cell lines (43 of 46 genes mapped between RNAseq and the CRISPR screen), comparing the treatment-resistant derivative (Gem-, Cis-, GemCis-resistant) to the associated parental cell line. Asterisks indicate a statistically significant result (moderated t-test, *FDR < 0.05). The bar plot on top is the aggregate count of significant results across all 15 comparisons. Genes are ranked by the count of statistically significant upregulated hits. (**B**) RNAseq (moderated t-test compared to parentals; *FDR < 0.05), (**C**) mass spectrometry proteomics (moderated t-test compared to parentals, *FDR < 0.25), and (**D**) CRISPR screen results for NPEPPS (mean ± SD; moderated t-test; *FDR < 0.05). (**E**) Representative immunoblots and densitometry quantification for independent triplicates (mean ± SEM) for NPEPPS in all cell lines (*FDR < 0.05).

#### NPEPPS drives cisplatin resistance *in vitro* and *in vivo*

To test our prioritization that NPEPPS regulates sensitivity to gemcitabine plus cisplatin treatment in GemCis-resistant BCa cells, and to parse its role in both cisplatin and gemcitabine resistance, we generated stable NPEPPS shRNA knockdowns in the KU1919-GemCis and T24-GemCis cell lines. We found that NPEPPS knockdown preferentially increased cisplatin, but not gemcitabine sensitivity (**Figure 3A, C**). Knockdown of NPEPPS did delay the growth of cells but did not have major effects on cell growth rates (**Figure S4A**). siRNA targeting of NPEPPS in the KU1919-GemCis cell line and shRNA and/or siRNA in T24-GemCis and 253J-GemCis cells confirmed our results that NPEPPS loss preferentially sensitizes cells to cisplatin (**Figure S4B,C**). Additionally, we used a gRNA from the CRISPR screen library to show that knockout of NPEPPS and the associated dose response matches our findings from shRNA and siRNA-mediated depletion of NPEPPS (**Figure S4D**). Conversely, overexpression of NPEPPS in KU1919 and T24 parental lines increased resistance to cisplatin, but not gemcitabine (**Figure 3B,D**). These results support NPEPPS as a regulator of sensitivity to gemcitabine plus cisplatin through its effect on regulating the cellular response to cisplatin.

**Figure 3.**
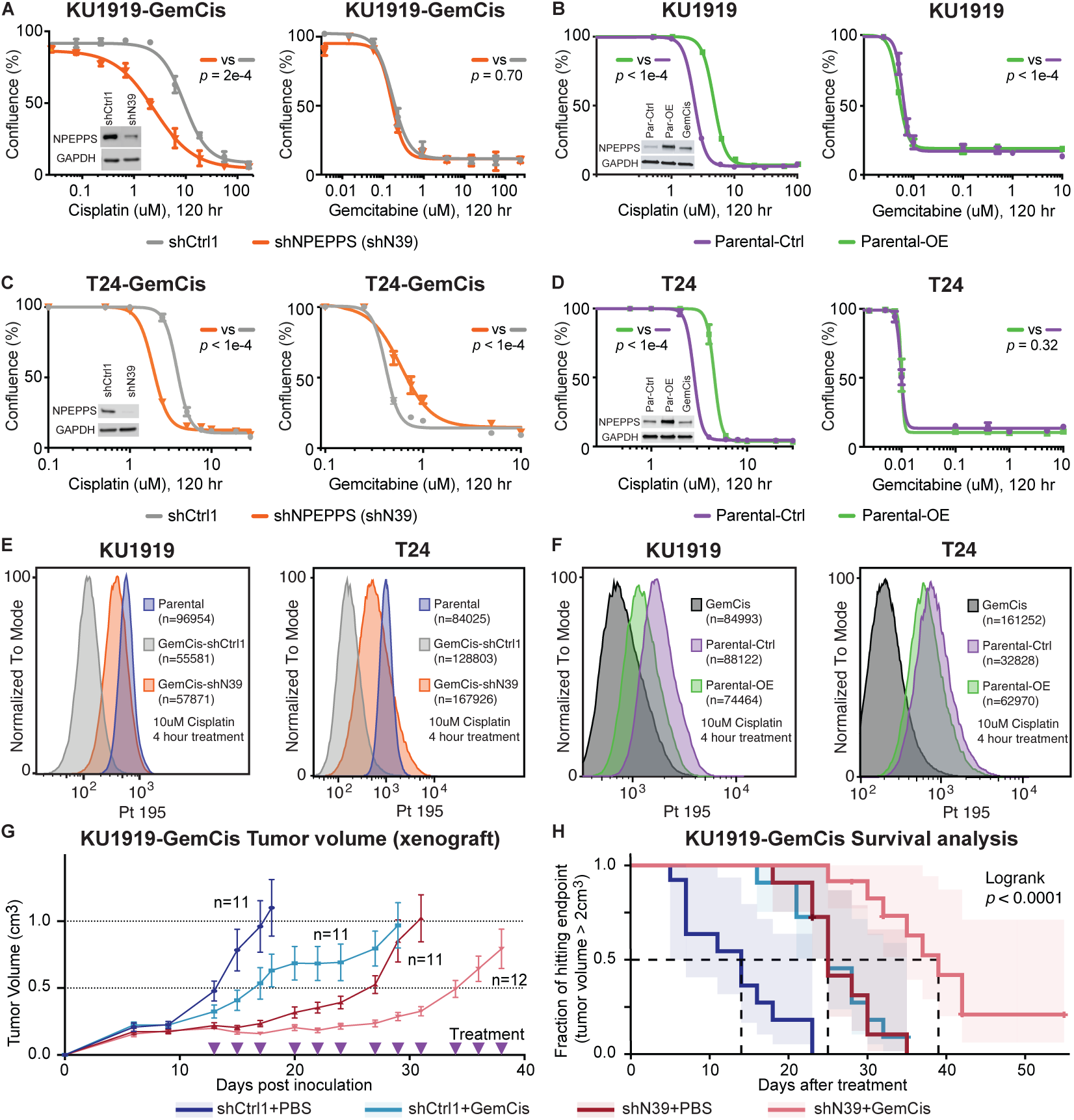
Genetic inhibition of NPEPPS resensitizes GemCis-resistant cells *in vitro* and *in vivo*. (**A, B**) KU1919-GemCis or T24-GemCis cells with knockdown of NPEPPS treated with increasing doses of cisplatin or gemcitabine. A total of 3 technical replicates per dose (mean ± SEM). (**C**, **D**). KU1919 or T24 parental cells with overexpression of NPEPPS treated with increasing doses of cisplatin or gemcitabine. A total of 3 technical replicates per dose (mean ± SEM). Independent experiments are reported in **Figure S4**. p-values comparing IC50 values using sum-of-squares F test. (**E,F**) Intracellular cisplatin levels in KU1919 T24 cells were measured after 4 hours of 10μM cisplatin treatment using CyTOF, with the number of live cells analyzed as indicated. (**G**) Tumor volume (mean ± SEM) of KU1919-GemCis xenografts measured over time and across 4 treatment groups considering non-targeting shRNA controls (shCtrl1), shRNA targeting NPEPPS (shN39), PBS vehicle control (PBS), or gemcitabine plus cisplatin treatment (GemCis). (**H**) Survival analysis of xenograft models with a defined endpoint of a tumor volume > 2cm^3^. Logrank test was applied to test significance.

To evaluate NPEPPS impact on intracellular cisplatin concentrations, we directly measured intracellular cisplatin using the metal ion detection capabilities of cytometry by time-of-flight, CyTOF (65). We measured intracellular cisplatin after 4 hours of treatment at 10μM in the same KU1919 and T24 cells evaluated for dose response. Using KU1919 as the illustrative example, KU1919-GemCis cells (median Pt 195 = 102) showed lower cisplatin concentration compared to KU1919 parental cells (median Pt 195 = 565). Control knockdown had little effect (median Pt 195 = 121), but NPEPPS knockdown shifted the intracellular levels of cisplatin to the parent lines (median Pt 195 = 375), suggesting that NPEPPS depletion increased intracellular cisplatin (**Figure 3E** and **S5**). These findings were replicated in the T24 cell lines (**Figure 3E** and **S5**). In the overexpression setting, and using KU1919 as the example, KU1919-GemCis cells (median Pt 195 = 763) again showed lower cisplatin concentration compared to KU1919 parental cells (median Pt 195 = 1706) with the overexpression control cells (median Pt 195 = 1738) showing little difference, but the NPEPPS overexpressing cells (median Pt 195 = 1203) showing a decrease in intracellular cisplatin (**Figure 3F** and **S6**). We found similar results with T24 cells (**Figure 3F** and **S6**). These data are consistent with the dose response results reported in **Figure 3A-D**.

To test if NPEPPS depletion sensitizes tumor cells to cisplatin-based chemotherapy *in vivo*, we established subcutaneous xenografts using the KU1919-GemCis cells with either NPEPPS shRNA knockdown or non-targeting shRNA control. When tumors reached roughly 200mm^3^, mice were randomized into four groups: shCtrl1 with PBS (n=11), shCtrl1 with gemcitabine plus cisplatin (n=11), shN39 with PBS (n=11), and shN39 with gemcitabine plus cisplatin (n=12). Treatment was delivered through intraperitoneal injection, with PBS or gemcitabine plus cisplatin administered three times weekly for four weeks. Tumor volumes were monitored until they reached the predetermined endpoint of 2cm^3^. NPEPPS knockdown alone and gemcitabine plus cisplatin treatment alone had a significant impact on tumor growth compared to vehicle-treated, shRNA controls. The combination of NPEPPS knockdown and gemcitabine plus cisplatin treatment led to a stronger and more significant impact on tumor growth (**Figure 3G**). We further analyzed tumor growth using linear mixed-effects models aimed at capturing trends in tumor volume change in relation to pre-treatment baseline tumor volume across the four groups (**Figure S7A,B**). According to this model, tumor growth inhibition by NPEPPS knockdown (*p*=0.00178), GemCis treatment (*p*=5.49e-7), or the combination of NPEPPS knockdown and gemcitabine plus cisplatin treatment (*p*=1.47e-8) were all consistent effects over the treatment period (**Figure 3G,H**). We validated NPEPPS knockdown in the pre-xenograft inoculate cells and after tumors were removed from mice upon reaching the 2cm^3^ endpoint (**Figure S7C**). Survival analysis using tumor volume as the endpoint showed that mice treated with gemcitabine plus cisplatin had a 14-day survival advantage. Similarly, the knockdown of NPEPPS resulted in a 14-day survival advantage. Mice treated with gemcitabine plus cisplatin and with NPEPPS knockdown tumors had a 25-day survival advantage, a statistically significant improvement (Logrank test, p<0.0001) (**Figure 3H**).

#### NPEPPS regulates cisplatin response in MIBC patient-derived organoids

We next evaluated the role of NPEPPS in regulating cisplatin response in *ex vivo* expanded 3D primary cultures of MIBC tissue as patient-derived organoids (PDOs). PDOs are commonly used as an intermediate step to the clinical investigation of novel therapeutics as they more closely recapitulate the human tumor as supported by a high degree of clinical reliability and predictability of drug response in validation trials (66–69). As shown in **Figure 4A**, PDOs were generated from MIBC patients undergoing transurethral resection of a bladder tumor (T, n=3) or radical cystectomy (C, n=2). Three PDOs were generated prior to cisplatin-based chemotherapy (preChemo), one PDO was generated after patients received cisplatin-based chemotherapy (postChemo), and one PDO was generated from patients that did not receive cisplatin-based chemotherapy (noChemo). The tumor for patient 4 showed pathological complete response (pCR), while the tumors for the remaining chemotherapy-treated patients were pathological non-responders (pNR).

**Figure 4.**
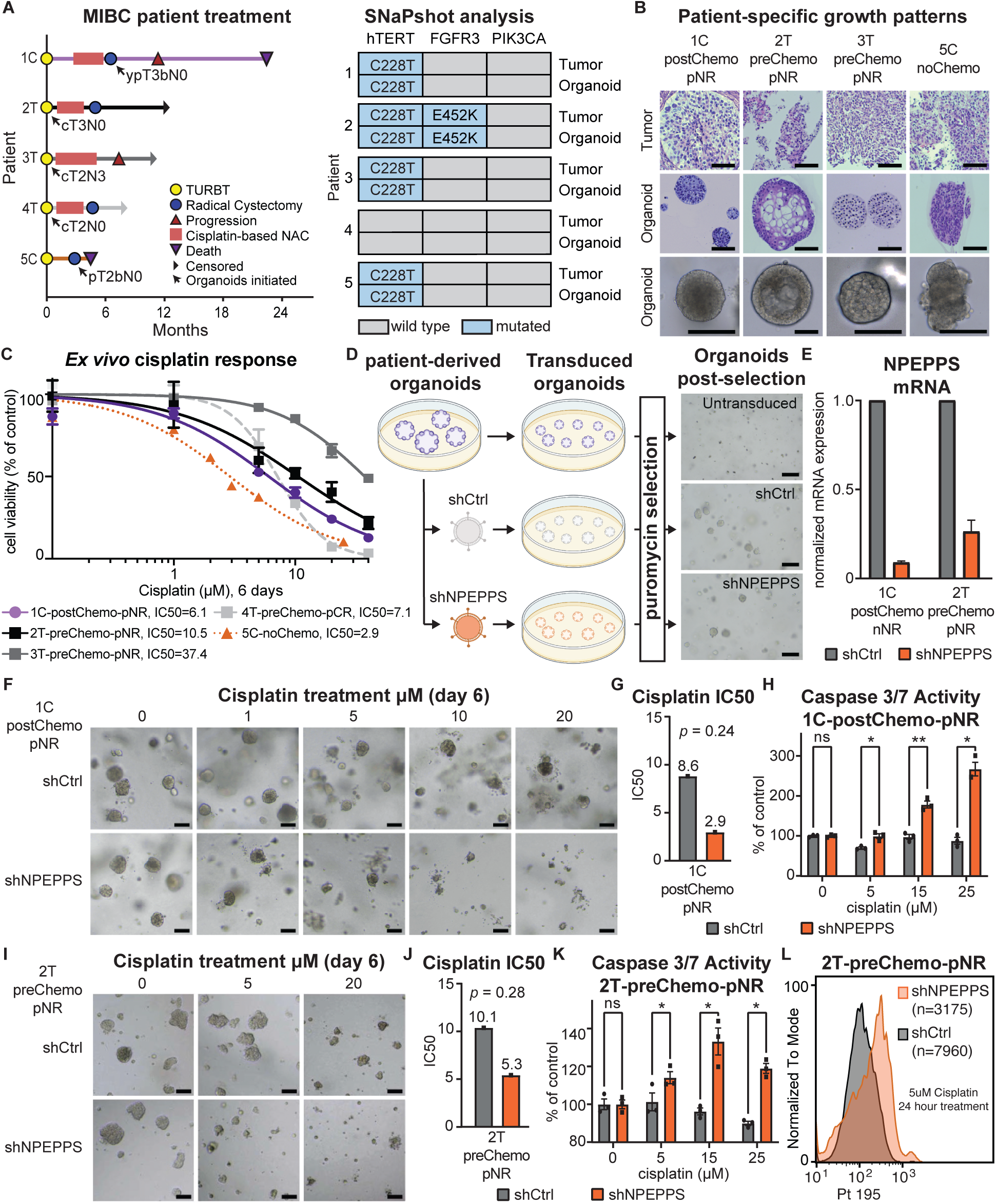
NPEPPS depletion sensitizes *ex vivo* models of bladder cancer to cisplatin. (**A**) Clinical course of muscle-invasive bladder cancer (MIBC) for patients including the point at which the patient tumor-derived organoid lines (PDO) were generated (indicated by the black arrow). Notation is the patient number followed by the tumor source, either T = TURBT = transurethral resection of bladder tumor, or C = radical cystectomy. (**B**) Representative bright-field images of organoids together with H&E staining of patient tumor-PDO pairs illustrating patient-specific growth patterns. Most PDOs exhibited round and dense structures, as represented by 1C-postChemo-pNR (Chemo = cisplatin-based chemotherapy regimen, pNR = pathological non-responder), although there was notable variation in growth and morphology. Scale bar = 100µm. (**C**) *Ex vivo* response of all nine PDOs treated with increasing concentrations of cisplatin. Cell viability, as a percentage of untreated control, was measured using CellTiter-Glo. The fitted dose-response curves represent viability corresponding to three biological replicate experiments and data are represented as mean ± SEM. pCR= pathological complete response. (**D**) Experimental workflow for lentiviral, shRNA-mediated NPEPPS-depletion in PDOs and representative images after puromycin selection. Scale bar = 400µm. (**E**) NPEPPS expression was evaluated by RT-PCR in shNPEPPS and shCtrl PDO lines normalized to cyclophilin. Error bars represent mean ± SD. (**F,I**) Representative bright-field images of control and NPEPPS-depleted PDOs, (**F**) 1C-postChemo-pNR or (**I**) 2T-preChemo-pNR,treated with the indicated cisplatin concentrations. Scale bar = 200µm. (**G,J**) IC50 values estimated from dose curves for cell viability measured through CellTiter-Glo (biological triplicates; mean ± SEM). (**H,K**) Relative caspase-3 and −7 activity in cisplatin-treated shCtrl and shNPEPPS PDOs. Caspase activity was measured by Caspase-Glo and normalized to untreated PDOs. (biological triplicates; mean ± SEM). (**L**) Intracellular cisplatin levels were measured after 24 hours of 5μM cisplatin treatment using CyTOF, with the number of live cells analyzed as indicated.

The tumor origin of the PDOs was confirmed by the detection of matching bladder cancer-specific mutations in tumor-organoid pairs (**Figure 4A**). As expected with this patient-derived platform, PDOs displayed distinct and tumor-specific phenotypes with notable variations in cell density and roundness (**Figure 4B**). We performed dose response with cisplatin to determine patient-specific *ex vivo* responses. PDOs displayed a range of responses to cisplatin, with a median cisplatin IC50 value of 6.1 µM, a minimum of 2.9 µM, and a maximum of 37.4 µM **(Figure 4C**).

To investigate the effect of NPEPPS depletion on cisplatin resistance, we selected two PDO lines from patients with clinical neoadjuvant chemotherapy (NAC)-resistance and intermediate *ex vivo* cisplatin resistance (1C-postChemo-pNR and 2T-preChemo-pNR), and performed shRNA-mediated knockdown of NPEPPS (**Figure 4D-E**) followed by their treatment with cisplatin for six days (**Figure 4F,I**). NPEPPS depletion lowered the cisplatin IC50 values by approximately 5µM in both of the tested PDOs (**Figure 4G,J**). Furthermore, significantly increased caspase-3 and −7 activity was observed following three days of cisplatin treatment in shRNA-mediated NPEPPS depletion compared to control PDOs, suggesting that NPEPPS depletion increases cisplatin-mediated apoptosis (**Figure 4H,K)**. Finally, we show that intracellular cisplatin levels in the shRNA control organoids (median Pt 195 = 113) are increased with the knockdown of NPEPPS (median Pt 195 = 213; **Figure 4L**). Collectively, these results demonstrate that genetic inhibition of NPEPPS sensitizes PDOs to cisplatin, which is consistent with our findings in the bladder cancer cell lines.

Next, we tested whether NPEPPS independently increases cisplatin resistance. We selected treatment-naïve (5C-noChemo) and post-NAC (1C-postChemo-pNR) PDO lines to represent treatment-naïve and cisplatin-treated patients with exogenously expressed NPEPPS via lentiviral transduction (**Figure 5A**). NPEPPS overexpression was confirmed by RT-qPCR, showing a 10-fold overexpression for 5C-noChemo and 3-fold overexpression for 1C-postChemo-pNR (**Figure 5B**). Cisplatin dose response was measured and we observed a decrease in cisplatin-mediated apoptotic blebbing and loss of structure in NPEPPS-overexpressing PDOs compared to empty vector control PDOs (**Figure 5C,G**). Furthermore, from the estimated IC50 values, we observed an increase in resistance to cisplatin associated with increased NPEPPS expression (**Figure 5D,H**). Moreover, less apoptosis was observed in NPEPPS-overexpressed PDOs treated with 15 µM and 25 µM cisplatin, as measured by caspase-3 and −7 (**Figure 5E,I**). Again, we found that NPEPPS controlled intracellular cisplatin concentrations, with the NPEPPS overexpressing cells (median Pt 195 = 682) showing decreased levels of cisplatin compared to the empty vector control cells (median Pt 195 = 996; **Figure 5F**). Taken together, these results suggest that NPEPPS regulates cisplatin response in treatment naïve and post-NAC treatment PDOs, highlighting NPEPPS as an attractive therapeutic target across cisplatin-based chemotherapy settings.

**Figure 5.**
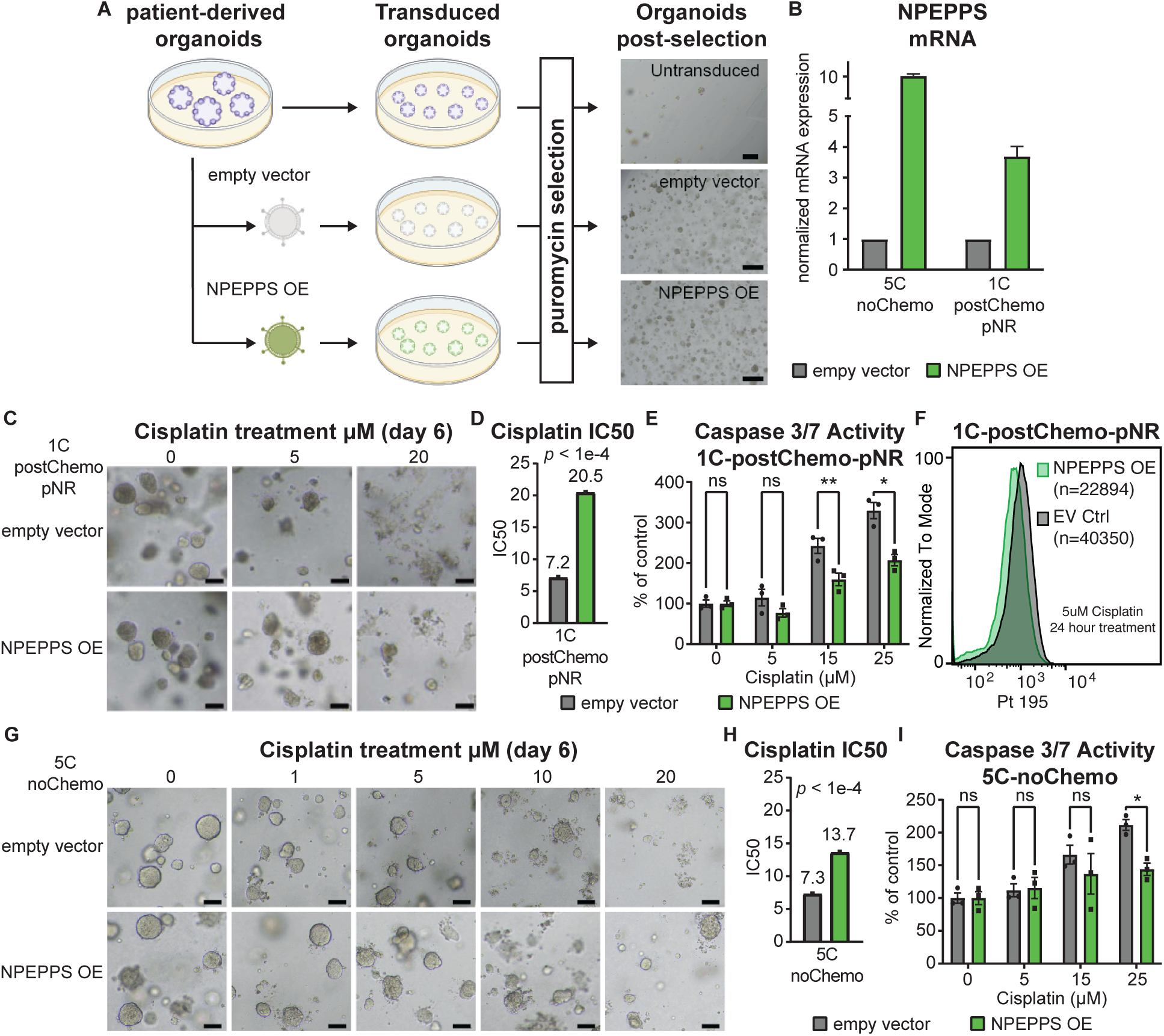
*Ex vivo* NPEPPS overexpression enhances cisplatin resistance. (**A**) Schematic representation of the experimental procedure for NPEPPS overexpression in PDOs and representative images after puromycin selection. Scale bar = 400µm. **B)** NPEPPS expression was evaluated by RT-PCR in shNPEPPS and shCtrl PDO lines normalized to cyclophillin. Error bars represent mean ± SD. (**C,G**) Representative bright-field images of empty vector control and NPEPPS-overexpressing PDOs, (**C**) 1C-postChemo-pNR or (**G**) 5C-noChemo, treated with the indicated cisplatin concentrations. Scale bar = 400µm. (**D,H**) IC50 values estimated from dose curves for cell viability measured through CellTiter-Glo (biological triplicates; mean ± SEM). (**E,I**) Relative caspase-3 and −7 activity in cisplatin-treated empty vector control and NPEPPS-overexpressing PDOs. Caspase activity was measured by Caspase-Glo and normalized to untreated PDOs. (biological triplicates; mean ± SEM). (**F**) Intracellular cisplatin levels were measured after 24 hours of 5μM cisplatin treatment using CyTOF, with the number of livecells analyzed as indicated.

### Pharmacological inhibition of NPEPPS improves cisplatin response in MIBC organoids

To assess the pharmacological efficacy of targeting NPEPPS, we tested cisplatin treatment combined with the NPEPPS inhibitor, tosedostat (70). We focused on the three preChemo, TURBT-derived PDOs to test the efficacy of cisplatin-based chemotherapy in combination with tosedostat in treatment naïve MIBC patients that will subsequently receive pre-operative chemotherapy. From our initial evaluation of cisplatin sensitivity (**Figure 4C**), we found the most sensitive PDO in this group to be 4T-preChemo-pCR (IC50=7.1μM), followed by 2T-preChemo-pNR (IC50=10.5μM), and the most resistant to be 3T-preChemo-pNR (IC50=37.4μM). This range of sensitivities matches with the clinical response of patient tumors, where patient 4 had a pathological complete response (ypT0N0), patient 2 had residual disease following NAC (ypT3aN0), and patient 3 progressed to metastatic disease during pre-operative chemotherapy (ypTxNxM+) (**Figure 6A**).

**Figure 6.**
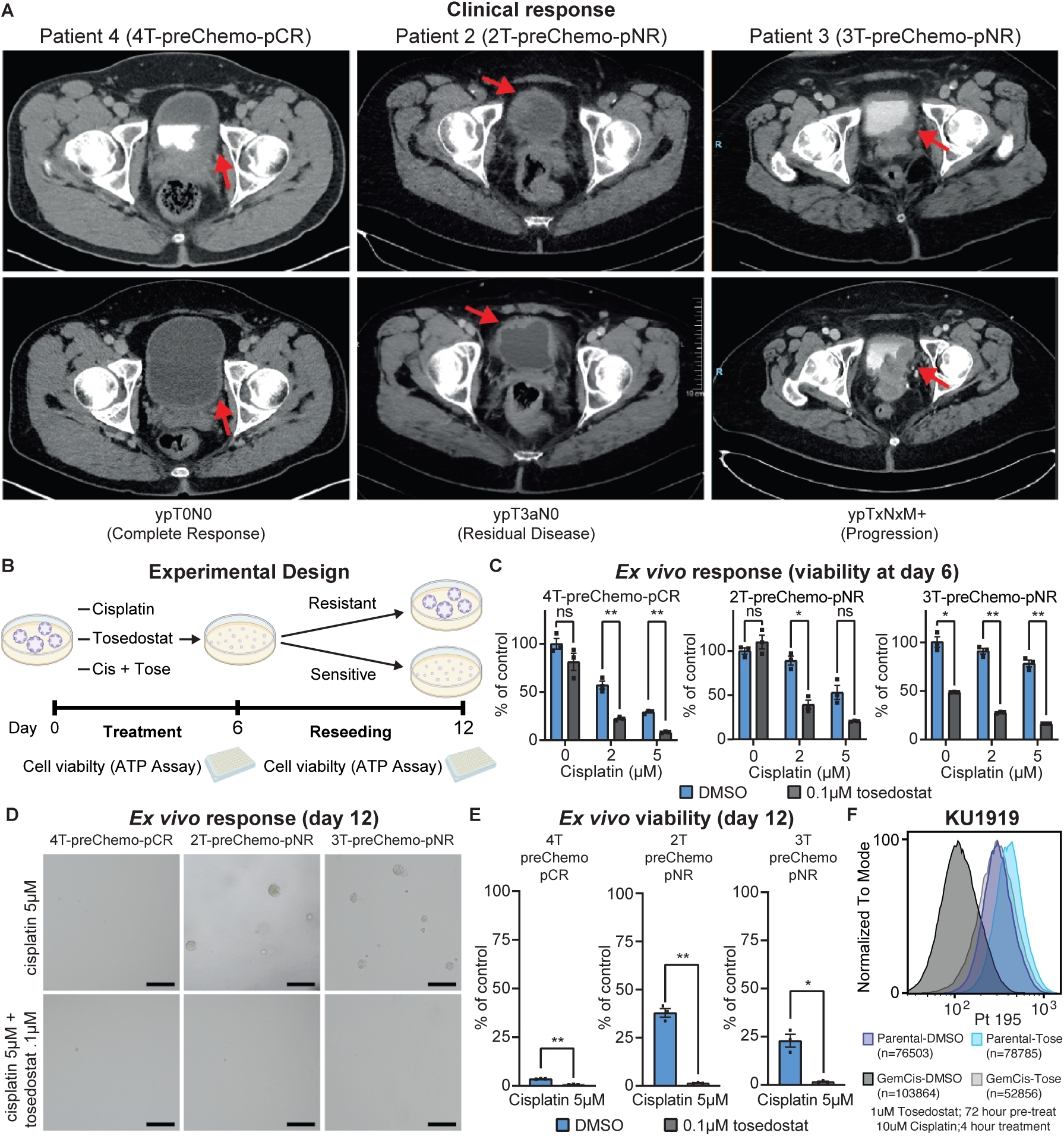
NPEPPS-inhibitor tosedostat overcomes cisplatin resistance ex vivo. (**A**) Clinical response to cisplatin-based chemotherapy in three MIBC patients. Pathological pre-operative chemotherapy response is annotated according to pathological stage following radical cystectomy or metastatic biopsy. Response is illustrated by pre-(top) and post-treatment (bottom) computerized tomography (CT) scans. Red arrows indicate bladder wall thickening and subsequence response. (**B**) Experimental design for treating PDOs with cisplatin and tosedostat alone or in combination. PDOs were withdrawn from treatment, dissociated to single cells and reseeded after 6 days. Cell viability was measured using CellTiter-Glo at day 6 and day 12. (**C**) *Ex vivo* response of PDOs treated with the indicated concentrations of cisplatin with or without the addition of tosedostat. After 6 days viability was measured by CellTiter-Glo. (biological triplicates; mean ± SEM). one-way ANOVA; *P<0.05, **P<0.005. (**D**) Representative bright-field images of reseeded organoids treated with cisplatin or cisplatin + tosedostat as indicated. Scale bar = 400µm. (**E**) Cisplatin response in reseeded organoids treated at the indicated concentrations of drug with viability measured by CellTiter-Glo. (biological triplicates; mean ± SEM). one-way ANOVA; *P<0.05, **P<0.005. (**F**) CyTOF results for KU1919-Parental or -GemCis cells treated with 1µM tosedostat or DMSO for 72 hours, followed by 10µM cisplatin for 4 hours.

Given that these PDOs are representative molecular (**Figure 4A**), morphological (**Figure 4B**), and pharmacological (**Figure 4C**) models of patient tumors, we treated the three PDOs with cisplatin, tosedostat, or the combination of cisplatin plus tosedostat (**Figure 6B**) based on previously identified plasma concentrations (70–75). Response to 2µM and 5µM cisplatin were consistent with our previous results (**Figure 4C**), with 4T-preChemo-pCR being the most sensitive to cisplatin (**Figure 6C, Figure S6A**). Tosedostat alone did not result in significant changes in viability for the 4T-preChemo-pCR and 2T-preChemo-pNR PDOs, but did result in a significant reduction in cell viability for 3T-preChemo-pNR, potentially highlighting a role for tosedostat as a monotherapy in some contexts (**Figure 6C, Figure S6A**). Significant reductions in cell viability were observed for all PDOs when cisplatin and tosedostat were combined after 6 days of treatment (**Figure 6C**, **Figure S6A**). Importantly, cisplatin response of 4T-preChemo-pCR was not compromised, but was further enhanced with the addition of tosedostat.

Next, we assessed the ability of drug-treated PDOs to re-establish, which is analogous to a clonogenic assay, and tests for cellular potential to outgrow following treatment (**Figure 6B,D**). After reseeding PDOs treated with 5μM cisplatin, 4T-preChemo-pCR showed an absence of organoid formation or outgrowth, while robust re-growth was observed for PDOs derived from tumors with residual or progressive disease (2T-preChemo-pNR and 3-preChemo-pNR) (**Figure 6D,E, Figure S6B**). Cisplatin plus tosedostat combination treatment eliminated PDO re-growth for all lines (**Figure 6E**). We additionally tested if tosedostat affected intracellular cisplatin concentrations similar to NPEPPS depletion. We found that priming bladder cancer cells with 1µM tosedostat for 72 hours followed by 10µM of cisplatin for 4 hours phenocopied NPEPPS depletion. Intracellular cisplatin in the KU1919-GemCis cells (median Pt 195 = 114) shifted to the same level as the KU1919-parental (media Pt 195 = 304) when pre-treated with tosedostat (median Pt 195 = 298). The parental lines also showed increased intracellular cisplatin after tosedostat pre-treatment (median Pt 195 = 410) (**Figure 3F**). Taken together, these findings provide strong evidence that tosedostat enhances cisplatin activity in clinically relevant and translatable experimental models of human bladder cancer, most likely by regulating intracellular cisplatin concentrations.

## DISCUSSION

NPEPPS has been suggested to play a role in a range of cellular processes including promoting autophagy, regulating cell cycle progression, and antigen processing (76–79). The majority of what is known about NPEPPS has been from studies in the brain, where it targets the degradation of polyglutamine sequences and misfolded protein aggregates associated with a number of neurodegenerative diseases, including Alzheimer’s disease, Huntington’s disease, and Parkinson’s disease (77,80–83). As reported in gnomAD, NPEPPS is a highly conserved gene and is constrained based on several metrics of intolerance to genetic variation in the population (84). NPEPPS is also ubiquitously expressed across human tissues (85). However, despite these features, genetic modification in mice is tolerable, though mice are slower growing, more sickly, and sterile (79,86), and as we have shown from our CRISPR screen results and follow-up experiments, knockout is not essential for growth in bladder cancer cells (**Figure 2D, S4**).

This work is not without its limitations. NPEPPS could have effects on treatment response outside of platinum drugs. For example, NPEPPS is upregulated in the Gem-resistant cell lines (**Figure 2**), and while we show that genetic NPEPPS loss is specific to cisplatin response (**Figure 3, S4**), NPEPPS upregulation could be part of broader cellular stress responses. Further studies will be needed to test other NPEPPS-mediated mechanisms of stress response. We note that all of our data support a cell autonomous effect of NPEPPS. As we indicated above, NPEPPS has been linked to mechanisms of immune response and non-cell autonomous effects of NEPPS were not tested here. Finally, we show that NPEPPS changes intracellular platinum drug concentrations. The mechanism by which NPEPPS regulates these intracellular levels requires future studies.

Despite these limitations, all of our data suggest that NPEPPS is a viable therapeutic target. Broadly, aminopeptidases have been therapeutically targeted as potential cancer treatments (87). More specifically, NPEPPS is a zinc-containing M1 aminopeptidase. Tosedostat was developed as a target of M1 aminopeptidases and the intracellular metabolized product CHR-79888 is the most potent inhibitor of NPEPPS reported (70,88). There have been a total of 11 clinical trials with tosedostat as reported in *clinicaltrials.gov* (73,74,88–90). The focus of its application has been in leukemias and myelomas, with several applications in solid tumors. The few clinical trials completed have reported tosedostat as being well tolerated by patients, but with modest effect as a single agent cancer treatment. A few examples of tosedostat in combination with cytarabine, azacitidine, capecitabine or paclitaxel have been tried, but there are no reports of tosedostat being tested in combination with platinum-based chemotherapy. This idea can also extend to the many patients that are ineligible for cisplatin-based chemotherapies by allowing a cisplatin dose reduction in the context of a cisplatin plus tosedostat combination. Furthermore, future determination of whether NPEPPS inhibition enhances carboplatin sensitivity may offer these patients an even more effective and less toxic drug combination option.

In conclusion, our finding that NPEPPS regulates cisplatin-based chemoresistance is both novel and actionable. Patient tumor-derived organoids (PDOs) have shown predictive value supporting personalized medicine in several tumor types (66–69,91,92). Our data with PDOs provides a strong preclinical rationale for the development of clinical trials evaluating tosedostat in combination with cisplatin (2).

## Acknowledgements

We would like to thank Megan Tu, Colin Sempeck, Ana Chauca-Diaz, Jason Duex, and Charles Owens for their help throughout this project. We would also like to thank Dania Manalo-Mae and the Cedars-Sinai Proteomics and Metabolomics Core Facility for the technical handling of the proteomic experiments.

