## Supplementary Figures for "NPEPPS is a novel and druggable driver of platinum resistance"

### Supplementary Figures and Tables for
#### NPEPPS is a novel and druggable driver of platinum resistance


Robert T. Jones[1,14], Mathijs Scholtes[2,14], Andrew Goodspeed[1,3,14], Maryam Akbarzadeh[2,5], Saswat Mohapatra[4], Lily Elizabeth Feldman[1], Hedvig Vekony[1], Annie Jean[1], Charlene B. Tilton[1], Michael V. Orman[1], Shahla Romal[5], Cailin Deiter[1], Tsung Wai Kan[2,6], Nathaniel Xander[1], Stephanie Araki[1], Molishree Joshi[1,7], Mahmood Javaid[8], Ryan Layer[8,9], Teemu D. Laajala[1,10], Sarah Parker[11], Tokameh Mahmoudi[2,5,6*], Tahlita Zuiverloon[2,*], Dan Theodorescu[4,12,13,*], and James C. Costello[1,3,15,*]

[1]Department of Pharmacology, University of Colorado Anschutz Medical Campus, Aurora, CO, USA
[2]Department of Urology, Erasmus MC Cancer Institute, Erasmus University Medical Center Rotterdam, Rotterdam, The Netherlands
[3]University of Colorado Cancer Center, University of Colorado Anschutz Medical Campus, Aurora, CO, USA
[4]Cedars-Sinai Samuel Oschin Comprehensive Cancer Institute, Los Angeles, CA, USA
[5]Department of Biochemistry, Erasmus University Medical Center Rotterdam, Rotterdam, The Netherlands
[6]Department of Pathology, Erasmus University Medical Center Rotterdam, Rotterdam, The Netherlands
[7]Functional Genomics Facility, University of Colorado Anschutz Medical Campus, Aurora, CO, USA
[8]Computer Science Department, University of Colorado, Boulder
[9]BioFrontiers Institute, University of Colorado, Boulder
[10]Department of Mathematics and Statistics, University of Turku, Turku, Finland.
[11]Smidt Heart Institute & Advanced Clinical Biosystems Research Institute, Cedars Sinai Medical Center, Los Angeles, California 90048, United States
[12]Department of Urology, Cedars-Sinai Medical Center, Los Angeles, CA, USA
[13]Department of Pathology and Laboratory Medicine, Cedars-Sinai Medical Center, Los Angeles, CA, USA
[14]Equal first authors
[15]Lead Contact
*Corresponding Authors

**Corresponding Authors**

Tokameh Mahmoudi, PhD
Departments of Pathology, Urology and Biochemistry
Erasmus University Medical Center
Dr. Molewaterplein 40, 3015GD, Rotterdam
The Netherlands
+31 6 53152973


Tahlita Zuiverloon, MD, PhD
Department of Urology
Erasmus MC Cancer Institute, Erasmus University Medical Center

Dr. Molewaterplein 40, 3015GD, Rotterdam
The Netherlands
+31 6 26 41 90 87


Dan Theodorescu, MD, PhD
Departments of Surgery and Pathology
Cedars-Sinai Medical Center
8700 Beverly Blvd. OCC Mezz C2002
7/3/23 10:07:00 AMLos Angeles, CA 90048
+1 (310) 423-8431


James C Costello, PhD
Department of Pharmacology
University of Colorado Anschutz Medical Campus
Mail Stop 8303
12801 E. 17th Ave., Rm L18-6114
Aurora, CO 80045
+1 (303) 724-8619


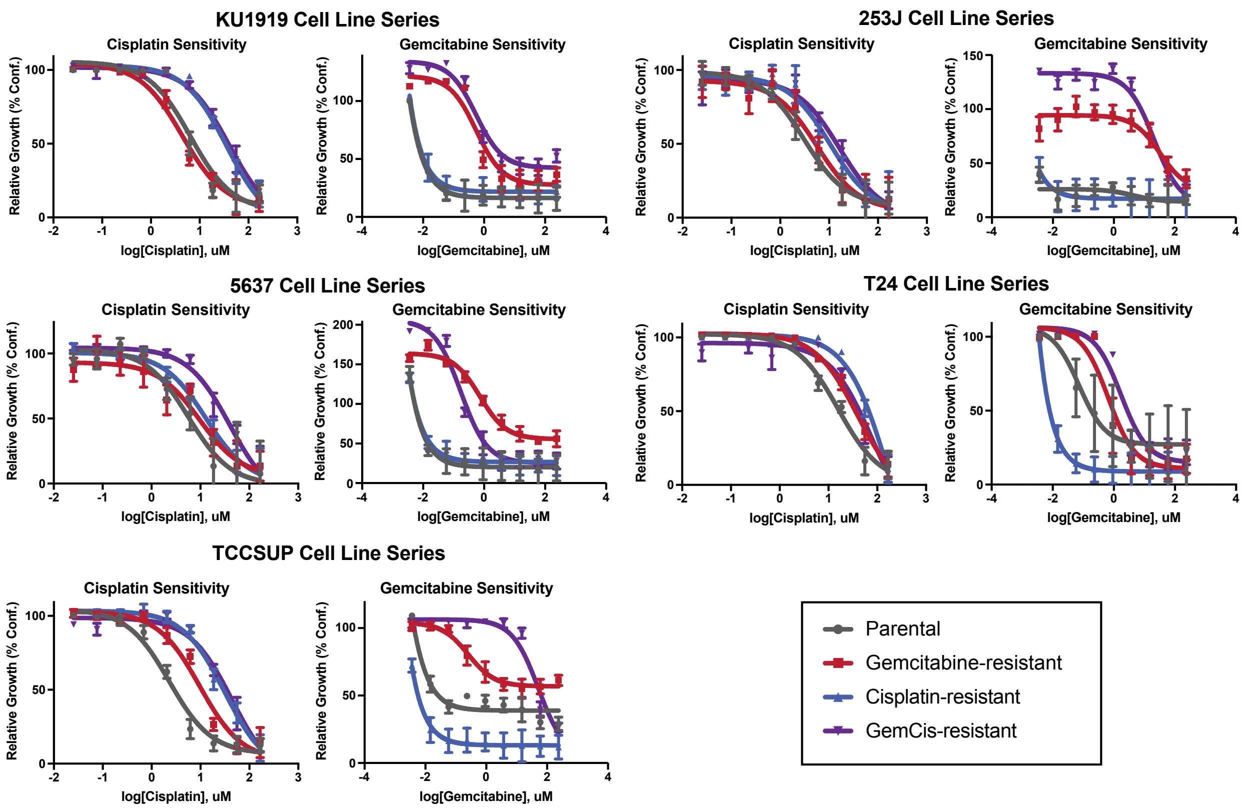


**Figure S1. Dose response for all cell line derivatives.** Parental, cisplatin-resistant, gemcitabine-resistant, and gemcitabine plus cisplatin (GemCis)-resistant cells for each of the five bladder cancer cell lines were treated with increasing doses of cisplatin or gemcitabine. Dose response curves were calculated. The resistant derivative lines were more resistant to the associated drug. Data represent a single experiment with each condition measured in technical triplicate wells (mean ± SEM).

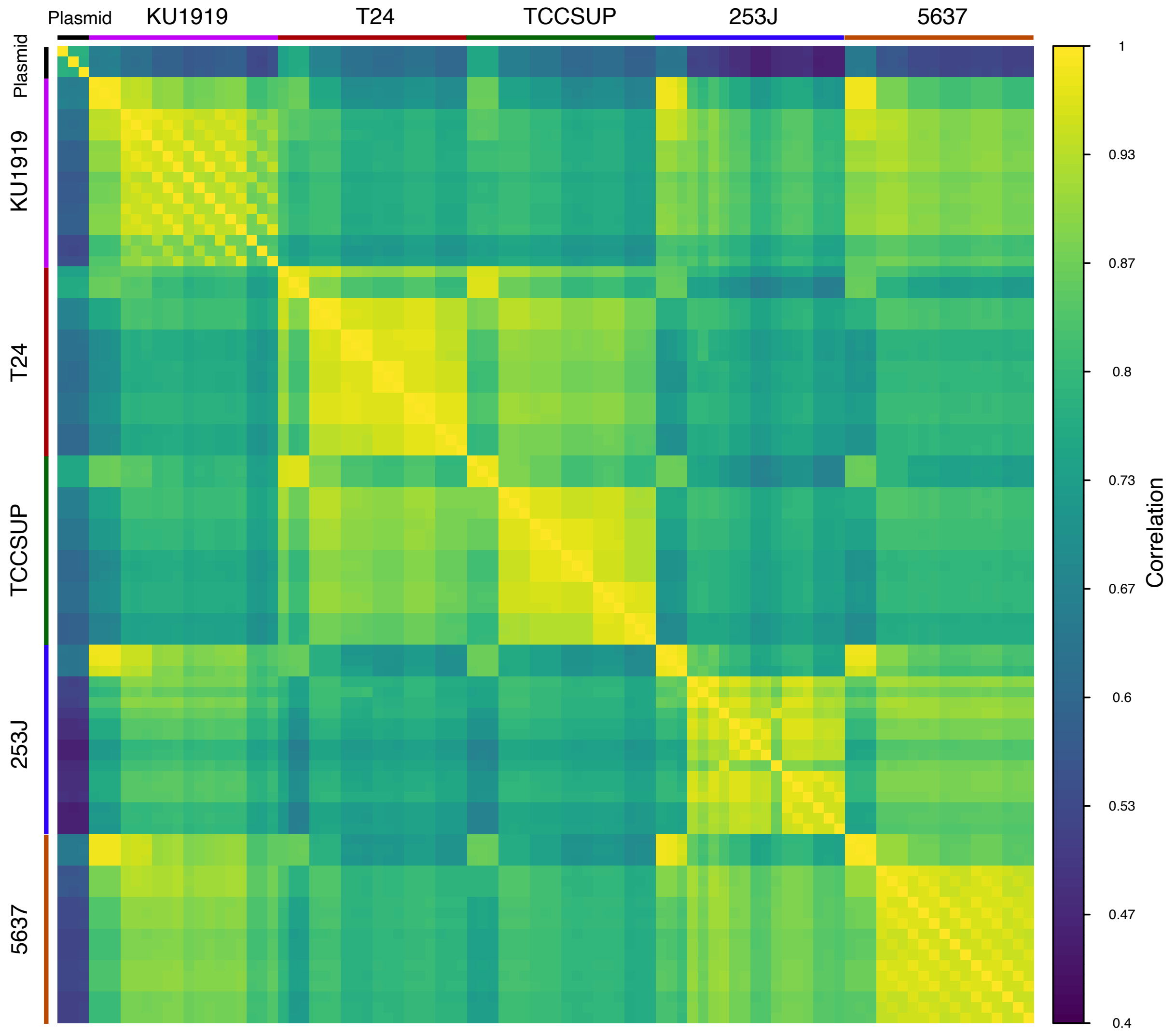


**Figure S2. Correlation of genes across conditions and cell lines from the CRISPR screen.** Pearson correlation was calculated between normalized gene counts across all tested conditions and replicates, from the original plasmid pool in triplicate to all cell lines. The ordering for each cell line includes: Day 0 (replicate 1, replicate 2, replicate 3), Day 10 (all replicates), Day 19 with PBS treatment (all replicates), Day 25 with PBS treatment (all replicates), Day 19 with gemcitabine plus cisplatin treatment (all replicates), and Day 25 with gemcitabine plus cisplatin treatment (all replicates).

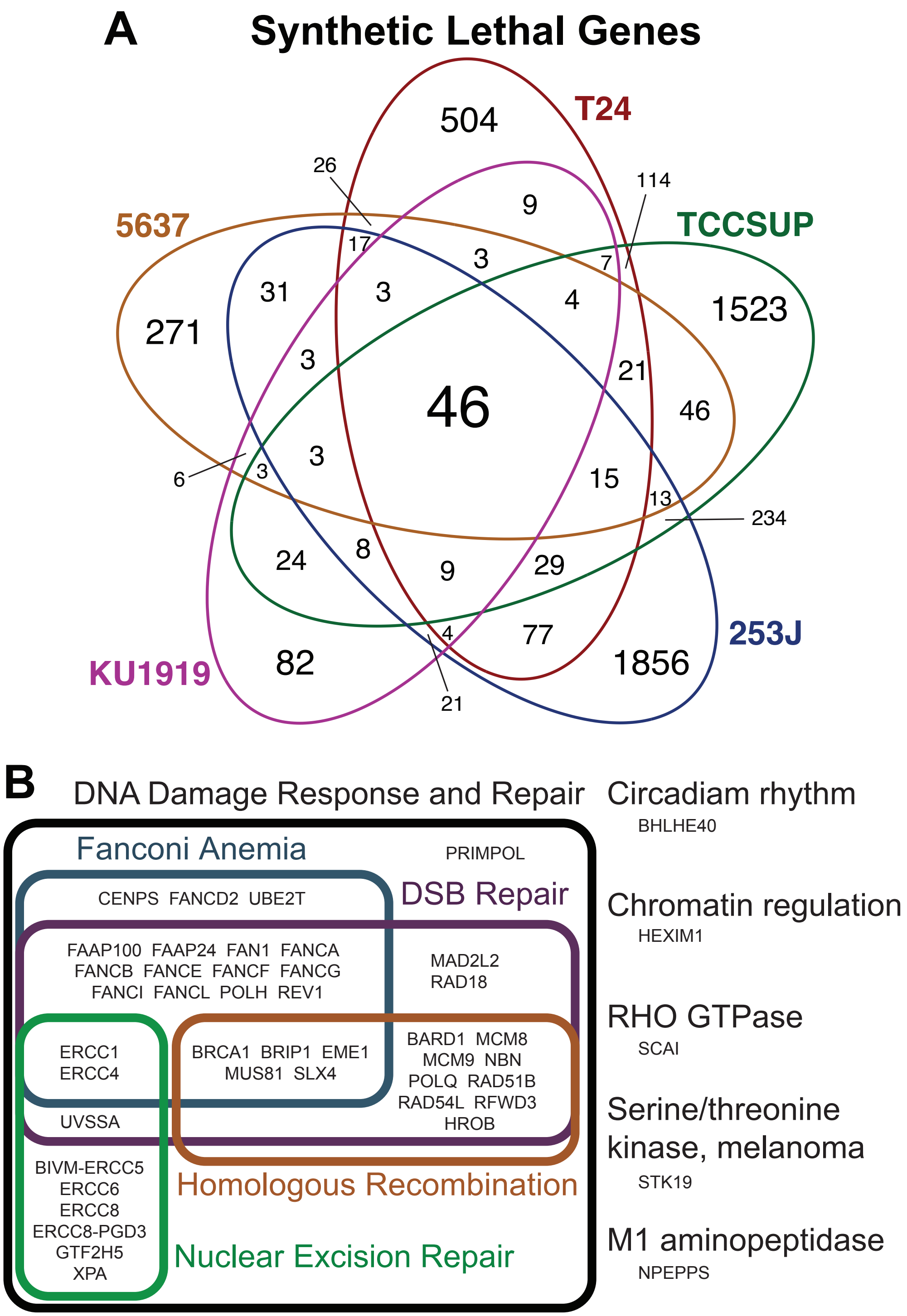


**Figure S3. Summary of synthetic lethal genes.** (**A**) Venn diagram of all gene counts across all 5 synthetic lethal screen results. The number of genes that were statistically significantly synthetic lethal (FDR < 0.05) are reported. (**B**) The putative annotations for the 46 commonly synthetic lethal genes. Detailed annotation information can be found in **Table S7**.

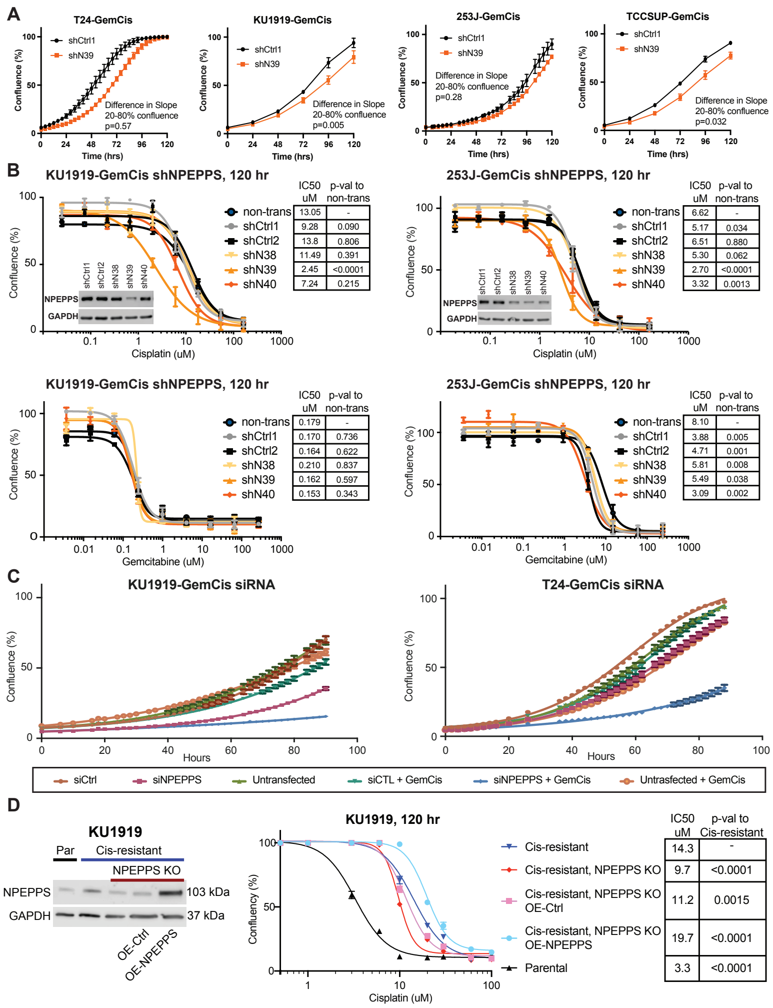


**Figure S4. shRNA and siRNA targeting NPEPPS resensitizes to cisplatin and gemcitabine plus cisplatin.** (**A**) Growth of GemCis-resistant cells with control knockdown (shCtrl1) or knockdown of NPEPPS (shN39) were measured over 120 hours in control (PBS) treatment conditions. Growth rates were calculated as the slope of the curve between 20%-80% confluency. (**B**) Dose response for shRNA targeting NPEPPS (shN38 = TRCN0000073838; shN39 = TRCN0000073839; shN40 = TRCN0000073840) or non-targeting controls (shCtrl1 = MISSION pLKO.1-puro Non-Mammalian shRNA Control; shCtrl2 = MISSION pLKO.1-puro Non-Target shRNA Control). Cells were treated with cisplatin or gemcitabine separately. Data for KU1919-GemCis shown from two independent experiments with 3 technical replicates per dose (mean ± SEM). Data from 253J-GemCis represent a single experiment with 3 technical replicate wells per dose (mean ± SEM). Immunoblot analysis of NPEPPS protein across the different shRNAs and cell lines are inset in the cisplatin treatment graph. (**C**) Cell confluency was measured using Incucyte Zoom across untransfected cells, siRNA controls (siCtrl), and siRNA transfected cells targeting NPEPPS. Cells were treated with PBS or gemcitabine plus cisplatin (GemCis) at the resistant doses for each cell line (**Table S1**). Data shown represent a single experiment with 3 technical replicates (mean ± SEM) per timepoint represented. (**D**) Cisplatin-resistant KU1919 cells were CRISPR edited using a gRNA from the Brunello library, GCAAAGGCTGTAGTTGATGG. Overexpression of NPEPPS, along with the empty vector control (OE-Ctrl), was performed in the NPEPPS knockout cells. Immunoblot for NPEPPS protein expression is shown.

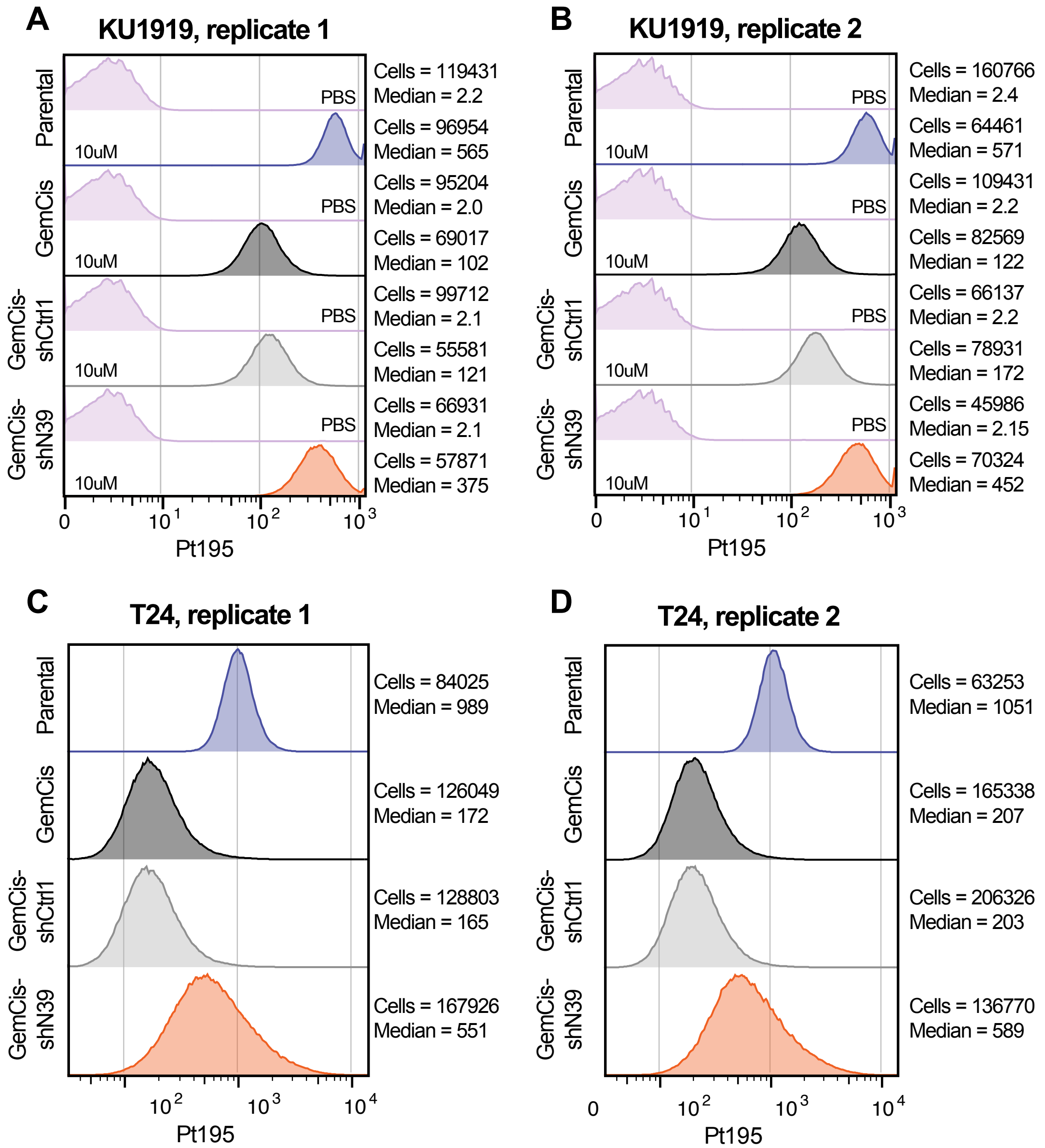


**Figure S5. Intracellular cisplatin measured using CyTOF.** (**A**) KU1919-Parental, KU1919-GemCis, KU1919-GemCis-shCtrl1, and KU1919-GemCis-shN39 cells were treated with PBS vehicle (0μM) or 10μM cisplatin for 4 hours and then intracellular cisplatin was measured using cytometry by time of flight (CyTOF). Results from replicates are shown. (**B**) The same experiment with 10μM cisplatin was performed with T24 cell lines and results from replicates are shown.

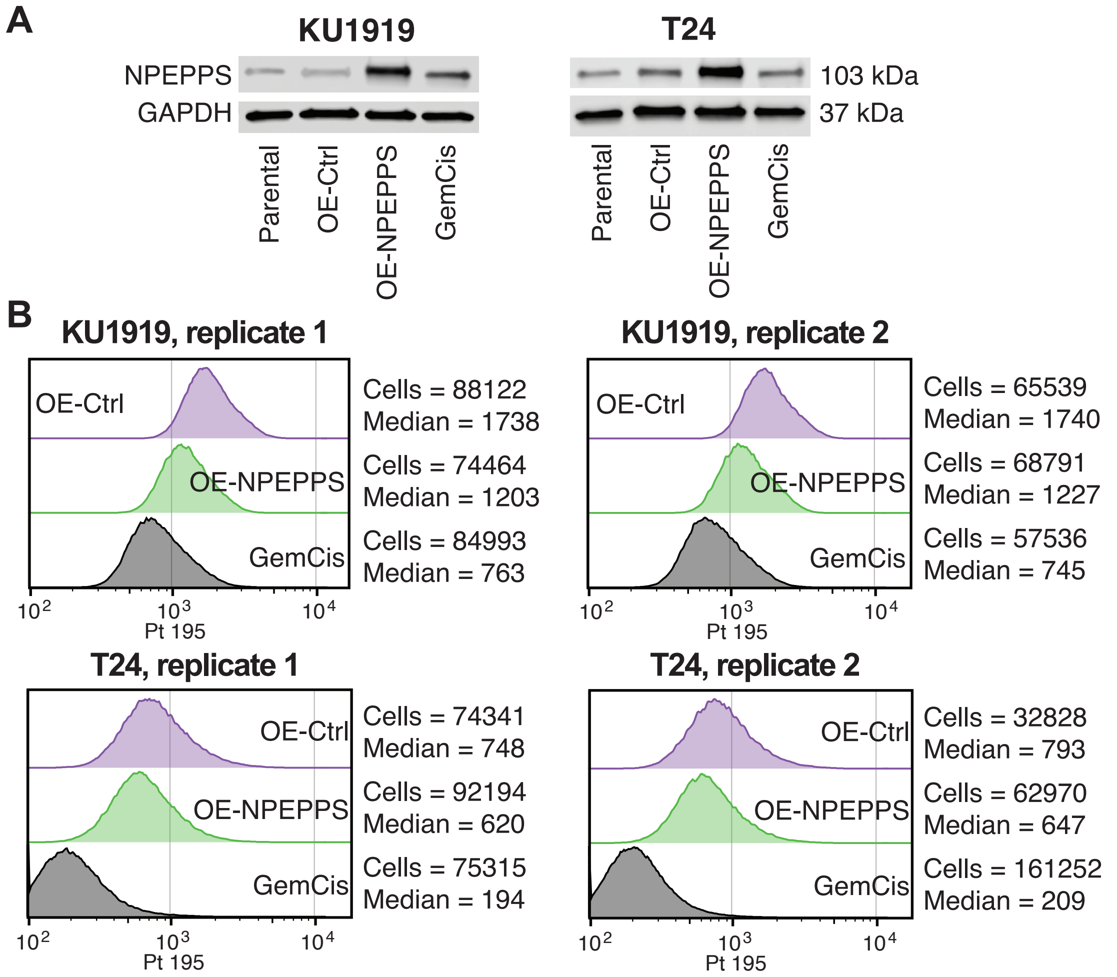


**Figure S6. Intracellular cisplatin in response to overexpression of NPEPPS.** (**A**) Immunoblot of NPEPPS to confirm the overexpression of NPEPPS. (**B**) CyTOF on KU1919 and T24 parental, overexpression control (OE-Ctrl), NPEPPS overexpression (OE-NPEPPS), or GemCis cells was used to measure intracellular cisplatin. The number of cells measured and the median value for each group are reported.

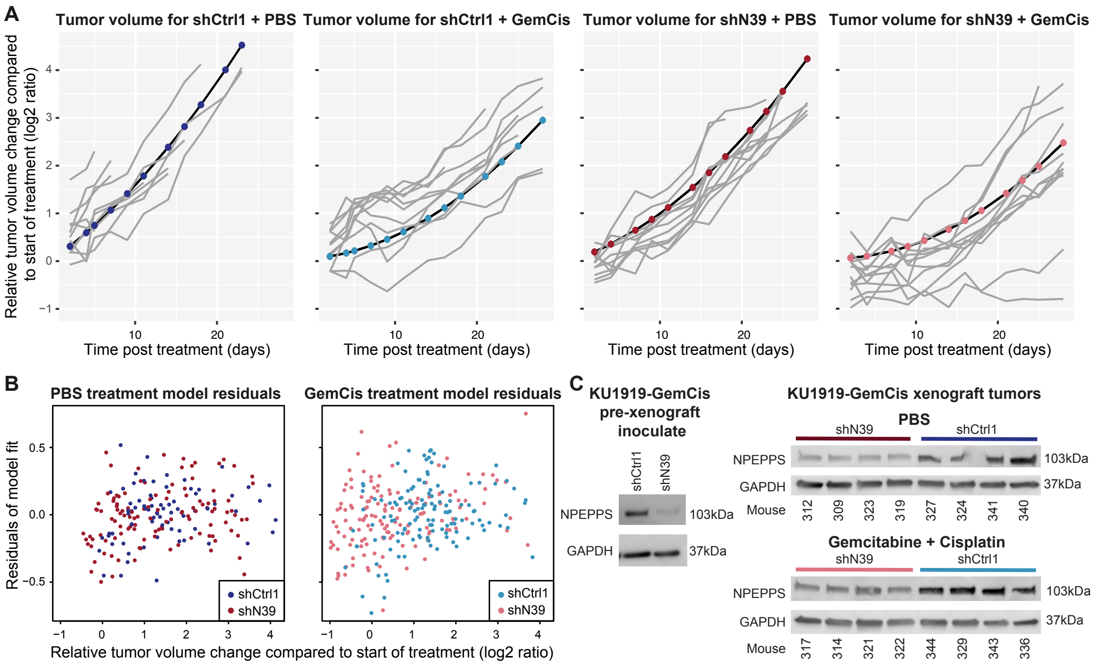


**D**

logrank test, p-values

| | | | |
|---|---|---|---|
| shCtrl1+GemCis | 7e-5 | | |
| shN39+PBS | 3e-5 | 0.8 | |
| shN39+GemCis | 4e-7 | 8e-4 | 2e-4 |
| | shCtrl1+PBS | shCtrl1+GemCis | shN39+PBS |

**Figure S7. KU1919-GemCis xenograft tumor growth modeling and validation.** (**A**) Fixed effects population-level model fit (thick lines) overlaid on top of the observations (grey lines). Longitudinal tumor volumes were divided by the baseline tumor volume and then $\log_2$-transformed before modelling. (**B**) Residuals for the final mixed-effects model (y-axis) coupled with the original observations (x-axis). No systematic trends were detected in model diagnostics, suggesting that the single fitted model successfully captured variation over the treatment arms and individuals. (**C**) Immunoblot on the left is from KU1919-GemCis cells that were injected into mice to establish tumors. Immunoblot on the right are from tumor samples after mice met the endpoint of the experiment of > $2cm^3$. (**D**) logrank test comparing all pairwise groups from **Figure 3G**.

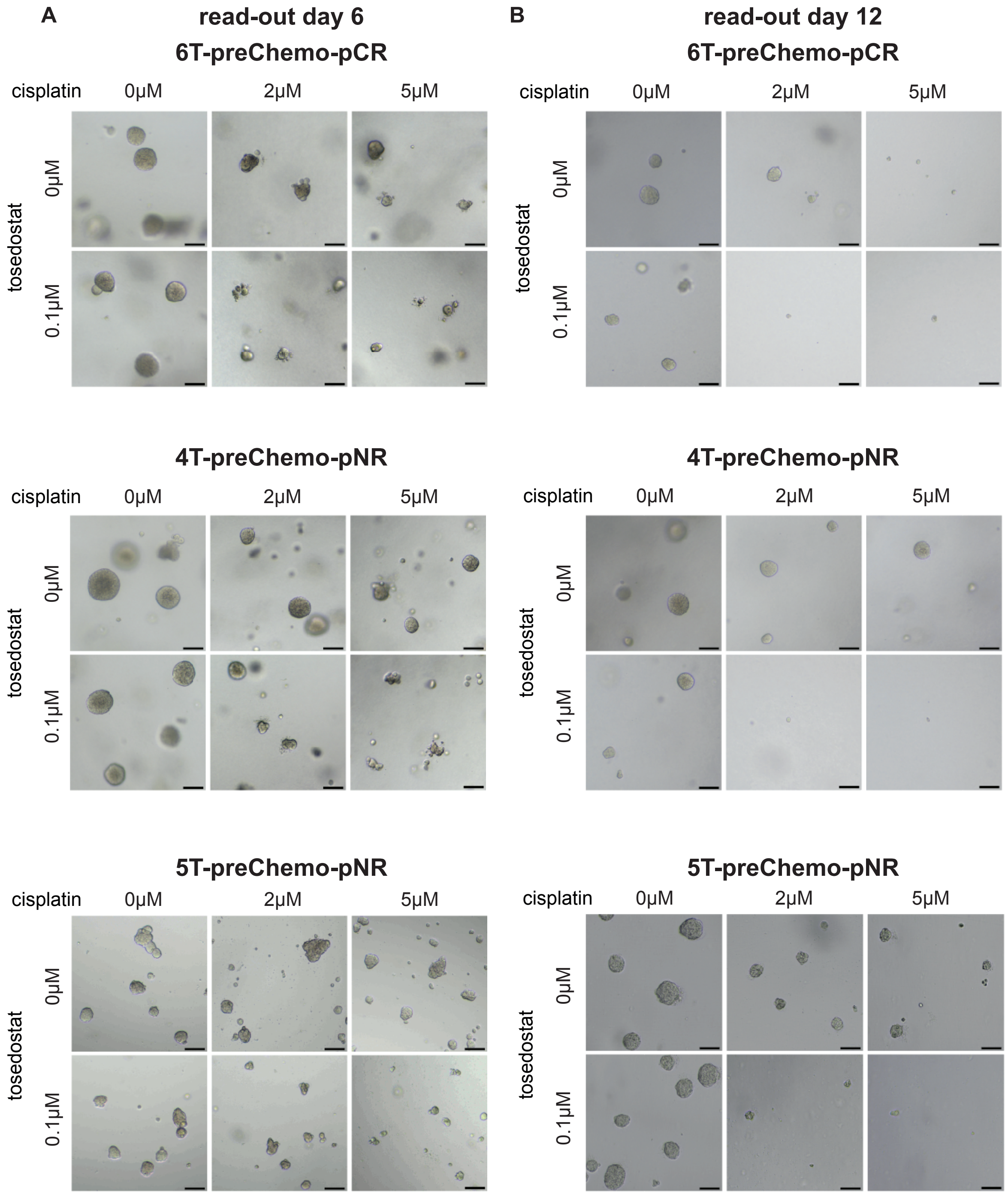


**Figure S6.** (**A**) Representative bright-field images of pre-POC organoids treated for six days at the indicated concentrations of drug. (**B**) Organoids originally treated at indicated concentrations of drug were dissociated to single cell and reseeded, allowed to grow new organoids for six days. Images were taken six days after reseeding, allowing organoids to re-grow. Scale bar = 200μm.

**SUPPLEMEMTAL TABLES**
Due to size, the supplemental tables can be downloaded here.

**Supplemental Table 1.** Established gemcitabine and cisplatin concentrations used for resistance cell line maintenance as reported by the RCCL. (https://research.kent.ac.uk/industrial-biotechnology-centre/the-resistant-cancer-cell-line-rccl-collection/)

**Supplemental Table 2**. Mutations filtered for quality and location in exonic gene regions.

**Supplemental Table 3**. Copy number alterations with the reported mean log2 ratio of the associated segment and the call of amplification (+), loss (-), or diploid (0).

**Supplemental Table 4.** CRISPR screening parameters used for each cell line.

**Supplemental Table 5.** Synthetic lethality as determined by gemcitabine plus cisplatin treated GemCis-resistant cells compared to saline treated cells with samples from day 19 and day 25 treated as covariates.

**Supplemental Table 6.** Gene Set Enrichment (fGSEA) results for synthetic lethal screen.

**Supplemental Table 7.** Gene annotations and references for top 46 hits from synthetic lethal screen.

**Supplemental Table 8**. Gene essentiality as determined by Day 25 saline treated samples compared to Day 0 samples in the GemCis-resistant cell lines.

**Supplemental Table 9.** Differential expression for RNAseq results for all derivatives compared to parental lines and for the KU1919, T24, TCCSUP, 253J, 5637, and all cell lines combined.

**Supplemental Table 10.** Differential expression from mass spec proteomics results for all derivatives compared to parental lines and for the KU1919, T24, TCCSUP, 253J, 5637, and all cell lines combined.

**Supplemental Table 11.** Alignment statistics from whole exome sequencing.

**Supplemental Table 12.** Comparison of mutation profiles reported through the CCLE.

**Supplemental Table 13.** Quality control measures for the CRISPR screening samples.

**Supplemental Table 14.** Broad Institute GPP LentiCRIPRv2 Library Prep Primers used in this study (P7 Index primer sequences A01-E09; P5 Staggered Pool).

**Supplemental Table 15.** Raw CRISPR screen counts.
